# A critical role for trkB signaling in the adult function of parvalbumin interneurons and prefrontal network dynamics

**DOI:** 10.1101/2020.06.28.175927

**Authors:** Nicolas Guyon, Leonardo Rakauskas Zacharias, Josina Anna van Lunteren, Jana Immenschuh, Janos Fuzik, Antje Märtin, Yang Xuan, Misha Zilberter, Hoseok Kim, Konstantinos Meletis, Cleiton Lopes-Aguiar, Marie Carlén

## Abstract

Inhibitory interneurons expressing parvalbumin (PV) in the prefrontal cortex (PFC) are central to excitatory/inhibitory (E/I) balance, generation of gamma oscillations, and cognition. Dysfunction of PV interneurons disrupts information processing and cognitive behavior. Tyrosine receptor kinase B (trkB) signaling is known to regulate the differentiation and maturation of cortical PV interneurons during development, but is also suggested to be involved in the activity and network functions of PV interneurons in the adult brain. Using a novel viral strategy for cell-type and region-specific expression of a dominant negative trkB in adult mice, we show that reduced trkB signaling in PV interneurons in the PFC leads to pronounced morphological, physiological, and behavioral changes. Our results provide evidence for a critical role of trkB signaling in the function of PV interneurons in the adult brain, local network activities central to prefrontal circuit dynamics, and cognitive behavior.

## Introduction

Tyrosine receptor kinase B (trkB) signaling elicited through its binding with brain-derived neurotrophic factor (BDNF) is crucial for a wide range of neurophysiological processes. Although trkB signaling was first characterized as instrumental in the development of the nervous system, it is now clear that this receptor has multiple roles in the adult nervous system, including the regulation of synaptic structure and plasticity, neurotransmitter release, cell morphology and excitatory-inhibitory dynamics in local circuits (Nagappan and Lu 2005; Park and Poo 2013; Kowiański et al. 2018). In addition to the full-length trkB (trkB.FL), C‐terminal truncated receptor (trkB.T) isoforms are expressed in different parts of the brain (Fenner 2012), with the trkB.T1 being expressed in both neurons and glia cells while the trkB.T2 and the trkB.Shc are expressed exclusively in neurons (Armanini et al. 1995; Stoilov, Castren, and Stamm 2002). The trkB.T isoforms hold the same extracellular and transmembrane domains as the full-length receptor, but their intracellular domain consists of a shorter sequence of amino acid residues that lacks the catalytic tyrosine kinase domain (Luberg et al. 2010). All trkB.T mediate BDNF-dependent signaling (Baxter et al. 1997; Wong and Garner 2012), however, the trkB.T1 is considered the prototype of truncated trkB receptors (Carim-Todd et al. 2009) and is thus the most studied truncated isoform (Fenner 2012). Mice with deletion of trkB.T1 do not display a severe developmental phenotype, which has led to the suggestion that trkB.T isoforms are involved in the regulation of BDNF, and in the function and differentiation of neurons rather than cell survival (Carim-Todd et al. 2009). In line with this, truncated receptors have been shown to function as a dominant-negative receptor inhibiting trkB signaling and BDNF function (Eide et al. 1996; Baxter et al. 1997; Stoilov, Castren, and Stamm 2002), but trkB.T1 is known to also hold BDNF independent functions, involved e.g., in the regulation of neuron and glia morphologies (reviewed in Fenner 2012). BDNF, and trkB signaling, have long been implicated in the etiology of mental disorders. Analysis of tissue from patients with schizophrenia has demonstrated a highly significant increase in trkB.T1 and trkB.Shc mRNA levels in the dorsolateral PFC, with a reduction of the trkB.FL/trkB.T ratio both on the mRNA and protein levels (Wong et al. 2013). Furthermore, decreased levels of prefrontal BDNF and trkB in schizophrenia have been correlated to decreased GAD67 mRNA expression, providing a link to aberrant GABA transmission (Hashimoto et al. 2005).

Neocortical GABAergic interneurons express trkB (Hashimoto et al. 2005), but not BDNF (Cellerino, Maffei, and Domenici 1996; Gorba and Wahle 1999). BDNF is secreted by excitatory neurons and contributes to the development of cortical interneurons, including through processes in which activity-regulated release of BDNF shapes the maturation of presynaptic cortical inhibitory synapses (Jiao et al. 2011; Huang et al. 1999). Much recent research has focused on trkB signaling in inhibitory interneurons expressing the calcium-binding protein parvalbumin (PV)(Zheng et al. 2011; Lucas, Jegarl, and Clem 2014; Xenos et al. 2018; Tan et al. 2018; Grech et al. 2019; Winkel et al. 2020). Cortical PV interneurons are densely interconnected (notably by gap-junctions) and distinguish themselves from other cortical inhibitory neuron types by their fast-spiking properties and the ability to sustain high firing frequencies (Tremblay, Lee, and Rudy 2016). PV interneurons regulate the excitability and spike timing of pyramidal neurons through both feedback and feedforward mechanisms (for review see Hu, Gan, and Jonas 2014). PV interneuron mediated inhibition, and the temporally locked interplay between inhibition and excitation, have been shown to be key to local network generation of oscillatory network activity in the gamma range (Cardin et al. 2009; Buzsáki and Wang 2012). Cortical gamma activity is associated with cognition, and altered gamma rhythms in the PFC are suggested to reflect the disease process in schizophrenia (Dienel and Lewis 2018). The activity of PV interneurons is also considered critical to the dynamic balance between excitatory and inhibitory neurotransmission (E/I balance) in the cortex (reviewed in Moore et al. 2010). Inhibition needs to flexibly respond to fluctuations in cortical state and balance the levels of excitatory input on the global activity level, while allowing for proper responses by distinct groups of excitatory neurons (Ferguson and Gao 2018; Sohal and Rubenstein 2019). The cortical PV interneurons are suggested to contribute to the formation of neuronal ensembles by restricting the ensemble size through suppression of less efficiently recruited neurons (Holtmaat and Caroni 2016). PV interneurons, thus, contribute to cortical function at several different scales, synaptic to network level, and it is only logical that this neuron subtype repetitively has been connected to etiology and/or pathophysiology in psychiatric disorders. Aberrant GABA transmission by PV interneurons inherently influences the E/I balance (Sohal and Rubenstein 2019), and the mechanisms altering the E/I balance have been intensively studied in order to shed light on the pathophysiology of psychiatric disorders. Studies of mouse models of schizophrenia and autism, as well as optogenetic experiments, have conclusively linked dysfunction of prefrontal PV interneurons to altered E/I ratio, deficits in information processing and cognitive behavior (Sohal et al. 2009; Yizhar et al. 2011; Carlén et al. 2012; Cho et al. 2015; Sohal and Rubenstein 2019).

Both trkB.FL and the truncated receptors are expressed in PV interneurons (Ohira and Hayashi 2009; Bracken and Turrigiano 2009). Studies of early postnatal genetic deletion of trkB in PV interneurons suggest that trkB signaling is critical to the function of this cell-type and the generation of gamma oscillatory synchrony (Zheng et al. 2011; Xenos et al. 2018). The data indicates that lack of trkB signaling in cortical PV interneurons increases the local network E/I ratio, negatively affecting cognition and behavior (Xenos et al. 2018; Tan et al. 2018). As these studies employ Cre/LoxP strategies for trkB deletion in transgenic mice, all trkB isoforms are deleted, and large parts of the brain are affected. Further, postnatal developmental processes involving BDNF/trkB signaling are impacted. Consequently, much regarding the role of trkB signaling in the adult brain is unknown. Furthermore, it has not been possible to conclusively pinpoint functional or behavioral alterations to region-specific changes in trkB signaling.

The aim of the current study was to investigate the role of BDNF/trkB signaling in the function of PV interneurons in the adult brain. The study focuses on the PFC as several lines of evidence strongly suggest a connection between aberrant trkB-mediated regulation of PV inhibition and prefrontal pathophysiology. For this we generated an adeno-associated viral (AAV) vector with Cre-dependent expression of a dominant negative trkB (trkB.DN), enabling spatially restricted targeting and temporally controlled expression onset. Adult expression of trkB.DN in inhibitory PV interneurons in the medial PFC (mPFC) led to pronounced morphological, physiological, and behavioral changes. The presented data demonstrate that BDNF/trkB signaling does not only regulate circuit development and plasticity of cortical PV interneurons but also their adult multidimensional functions in cortical dynamics and cognition.

## Results

### Targeting of trkB.DN to prefrontal PV interneurons in adult mice

To allow for spatially restricted and cell-type specific inhibition of trkB signaling in the adult brain, we generated an AAV vector with Cre-dependent expression of trkB.DN (AAV-DIO-trkB.DN-mCherry). Our trkB.DN construct holds intact extracellular and transmembrane domains, but the intracellular domain has been replaced by the monomeric red fluorescent protein mCherry (**Figs. 1a, S1a-c,** and **Methods)**. Thus, trkB.DN can compete for BDNF, but binding of BDNF to trkB.DN does not activate intracellular pathways. Further, as trkB.DN lacks any intracellular trkB domain, its expression does not induce any BDNF independent functions conveyed by truncated trkBs (Fenner 2012). In parallel we used single-nucleus RNA sequencing (snRNA-seq) (Märtin et al. 2019) to investigate the expression of *Ntrk2* (gene encoding for all trkB isoforms) and *Bdnf* in *Pvalb-*expressing interneurons in the adult mPFC. Vgat-expressing mPFC neurons from adult mice were isolated using FACS, and the molecular profile analyzed (n = 648 nuclei; **Fig. S1d**, and **Methods**). Clustering of gene expression data revealed 255 nuclei expressing *Pvalb*, with 253 nuclei co-expressing *Ntrk2*. Two clusters (14 and 15) expressed high levels of *Pvalb*, with 98.9% (173/175), and 100% (12/12) of the nuclei, respectively, also expressing *Ntrk2*; **Figs. S1d, e**). *Bdnf* was expressed in 2.9% (5/175) and 0% (0/12) of the nuclei in the two clusters (**Fig. S1e**). The data confirms the expression of *Ntrk2*, and the absence of autocrine mechanisms of BDNF, in prefrontal PV interneurons. The data further supports the notion that pyramidal neurons modulate GABAergic inhibitory activity of presynaptic interneurons through paracrine actions of BDNF (Jiao et al. 2011). For adult expression of trkB.DN in mPFC PV interneurons, the AAV was injected bilaterally (0.5 μl/injection site) into the mPFC of 8-12 weeks old PV-Cre mice (**Fig. 1b**). Control mice were generated by injection of an AAV with Cre-dependent expression of eYFP into PV-Cre mice (**Fig. 1b**). Throughout the current study, mice injected with AAV-DIO-trkB.DN-mCherry are referred to as trkB.DN mice, and mice injected with AAV-DIO-eYFP referred to as eYFP mice. Four weeks later, robust fluorescence was detected in prefrontal neurons in both groups of mice (**Fig. 1c**). Immunohistochemistry verified specific targeting of trkB.DN to mPFC PV interneurons (95.7 ± 3.1%; 1849/1940 trkB.DN-mCherry-expressing neurons also expressing PV; n = 3 trkB.DN mice; **Figs. 1d, e**). PV is a calcium-binding protein, with activity regulated expression, and the non-complete co-expression of PV and trkB.DN-mCherry is plausibly explained by PV expression below the detection threshold in a small subset of the PV interneurons (Sohal et al. 2009). The injected volume transduced 56.8 ± 4.1% (1849/3246) of the PV interneurons at the injection site (**Figs. 1d, e**).

**Figure 1.**
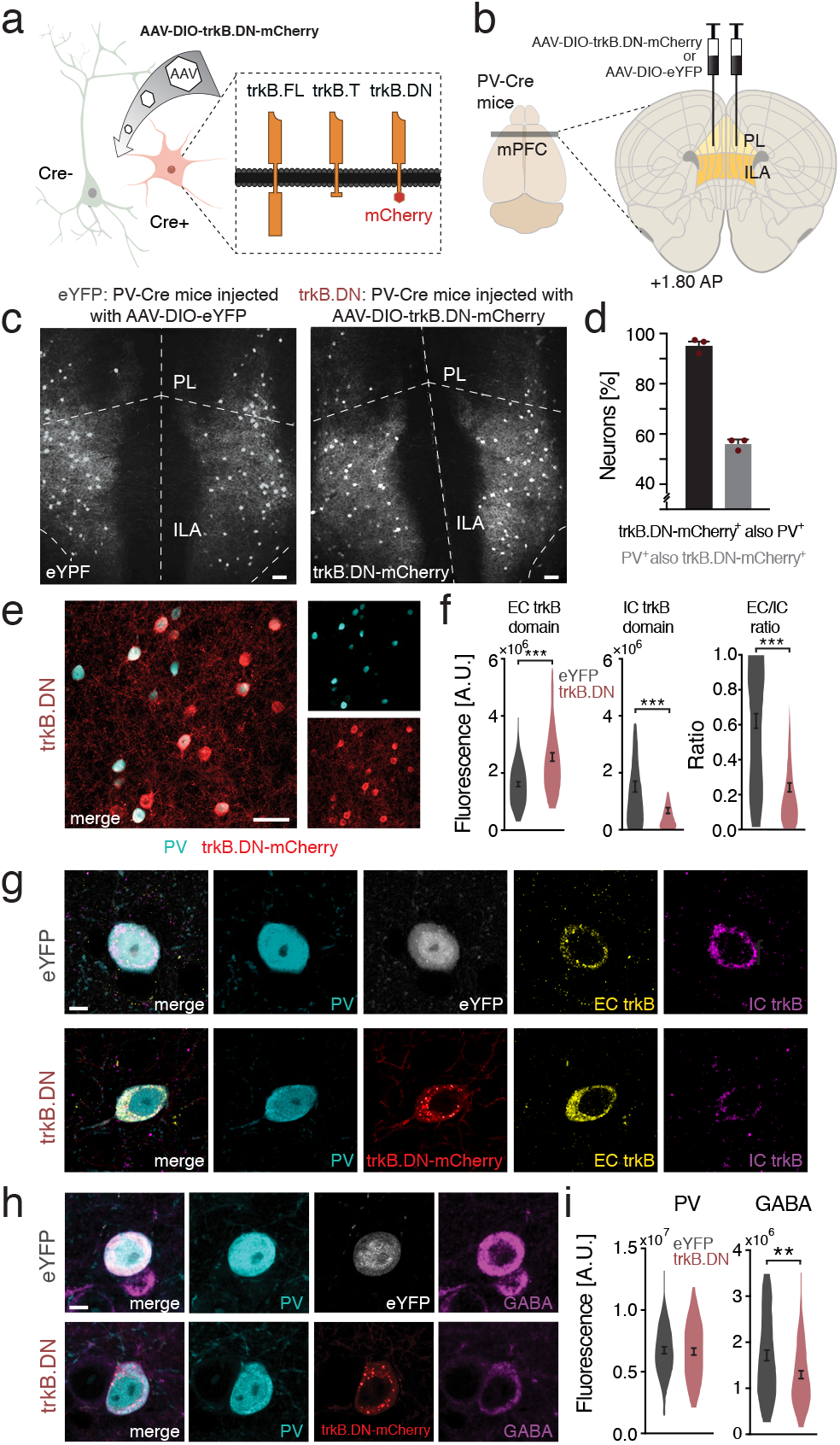
Targeting of trkB.DN to mPFC PV interneurons in adult mice. (**a**)Cre-expressing neurons transduced by AAV-DIO-trkB.DN-mCherry express the virally expressed trkB.DN-mCherry, and the endogenous full-length (trkB.FL) and truncated (trkB.T) trkB receptors. (**b**) AAV-DIO-trkB.DN-mCherry or AAV-DIO-eYFP was injected into the mPFC in adult (8-12 weeks old) PV-Cre mice, with the injection center targeted to the prelimbic area (PL). (**c**) Bilateral eYFP (left), and trkB.DN-mCherry (right), expression in mPFC neurons in PV-Cre mice four weeks after viral injections. PV-Cre mice were consistently used in the current study, and PV-Cre mice with AAV-DIO-trkB.DN-mCherry targeted to the mPFC are referred to as trkB.DN mice, and mice with AAV-DIO-eYFP targeted to the mPFC as eYFP mice. (**d**) Specificity and efficiency of targeting of trkB.DN to mPFC PV interneurons at the injection site (n = 3 trkB.DN mice). Black bar: percent trkB.DN-mCherry expressing neurons also expressing PV (95.7 ± 3.1%, 1849/1940 neurons); gray bar: percent mPFC PV interneurons also expressing trkB.DN-mCherry (56.8 ± 4.1%, 1849/3246 neurons). (**e**) Representative immunohistochemical detection of co-expression of PV (green) and trkB.DN-mCherry (red) in the mPFC of a PV-Cre mouse injected with AAV-DIO-trkB.DN-mCherry. (**f**) Levels of the extracellular (EC) trkB domain (left), the intracellular (IC) domain (middle), and the EC/IC ratio in mPFC PV interneurons expressing trkB.DN-mCherry (red; n = 99 neurons in 5 trkB.DN mice) or eYFP (gray; n = 74 neurons in 4 eYFP mice), respectively. EC domain: eYFP: 1613709 ± 85857 a.u.; trkB.DN-mCherry: 2571727 ± 145054 a.u., *p* < 0.0001. IC domain: 1522892 ± 191982 a.u.; trkB.DN-mcherry 675638 ± 103170 a.u., *p* < 0.0001. EC/IC ratio: 0.62 ± 0.04; trkB.DN-mcherry 0.25 ± 0.03, *p* < 0.0001. (**g**) Representative immunohistochemical detection of PV (cyan), EC trkB (yellow), IC trkB (pink), and eYFP (white) or trkB.DN-mCherry (red) in a mPFC PV interneuron in an eYFP mouse (top row), or a trkB.DN mouse (bottom row). (**h**) Representative immunohistochemical detection of PV (cyan), GABA (pink), and eYFP (white) or trkB.DN-mCherry (red), in a mPFC PV interneuron in an eYFP mouse (top row), or a trkB.DN mouse (bottom row). (**(i)** Left: The levels of PV protein in mPFC PV interneurons expressing eYFP (gray; 6757864 ± 276743 a.u.; n = 62 neurons from 3 eYFP mice) or trkB.DN-mCherry (red; 6641059 ± 284145 a.u.; n = 68 neurons from 3 trkB.DN mice), *p* = 0.7696. Right: The levels of GABA in mPFC PV interneurons expressing eYFP (gray; 1722901 ± 116046 a.u.) and trkB.DN-mCherry (red; 1298824 ± 85478 a.u.), *p* = 0.0077. Data shown as mean ± SEM. Unpaired test, Two-tailed was used to assess significance if data passed the D’Agostino & Pearson normality test, if not, the Wilcoxon rank-sum test was used. ** *p* < 0.01 *** *p* < 0.001. Scale bars: 75 μm in (c), 100 μm in (e), 10 μm in (g), 10 μm in (h).

To confirm the dominant negative action of trkB.DN we used immunohistochemistry to detect the extracellular domain of trkB in the cell bodies of virus-transduced PV interneurons in trkB.DN (n = 5), and eYFP (n = 4), mice (**Figs. 1f, g**). PV interneurons expressing trkB.DN expressed significantly higher levels of the extracellular trkB domain than PV interneurons expressing eYFP (**Fig. 1f)**, indicating an overall higher level of trkB in mPFC PV interneurons transduced by AAV-DIO-trkB.DN-mCherry. As mentioned, trkB.DN lacks the intracellular trkB domain. We therefore next investigated the levels of the intracellular domain and found that PV interneurons expressing trkB.DN expressed significantly lower levels of the intracellular trkB domain than PV interneurons expressing eYFP, reflecting a lower level of trkB.FL in the presence of virally expressed trkB.DN (**Fig. 1f)**. Concordantly, the extracellular/intracellular ratio was found to be significantly lower in mPFC PV interneurons expressing trkB.DN than in their eYFP expressing counterparts (**Fig. 1f**), and together these experiments demonstrate that trkB.DN acts as a dominant negative receptor. Immunohistochemistry revealed that adult expression of trkB.DN did not affect the levels of PV, but significantly reduced the levels of GABA in the cell bodies of mPFC PV interneurons (**Figs. 1h, i**). This contrasts findings of reduced PV levels in models of postnatal deletion of trkB, which affect the maturation of the PV neurons (Zheng et al. 2011).

### Morphological changes of trkB.DN expressing PV interneurons

For detailed study of cell morphologies and physiology, acute brain slices were prepared from cohorts of trkB.DN (n = 9), and eYFP (n = 15) mice. The detected fluorescent neurons were filled with biocytin and subjected to slice electrophysiology and/or detailed morphological analyses (**Figs. 2a, b**). The PV identity was confirmed in all investigated neurons (**Fig. 2b**). The cortical PV interneuron population contains basket neurons and a minor population of chandelier neurons (Tremblay, Lee, and Rudy 2016). Our reconstructions of biocytin-filled neurons identified neurons of both populations and revealed their strikingly differential morphology (**Fig. S2a**). However, due to the low sampling of chandelier neurons (trkB.DN mice: n = 4; eYFP mice: n = 7) only PV basket neurons (trkB.DN mice: n = 13; eYFP mice: n = 14) were included in our further morphological investigations. Reconstruction and three-dimensional (3D) morphometric analysis (Sholl analysis; **Fig. S2b**) of the biocytin-labeled neurons revealed an increased neurite (axons + dendrites) arborization in mPFC PV basket neurons expressing trkB.DN, particularly 100-200 μm from the soma (**Fig. 2c**). In addition, the total length of the neurites was significantly increased, as was the number of neurites (**Figs. 2d** and **S2d)**. However, the ending radius of the neurites was unchanged, indicating that the overall size of the neurons was unaltered (**Fig. S2d**). Both dendrites and axons displayed increased arborization and total length in mPFC PV interneurons expressing trkB.DN (**Figs. 2e-g**, and **S2e-m)**. Also, the number and the ending radius of the dendrites were significantly increased (**Fig. S2e).** However, the number of axons was not changed, nor the axonal ending radius (**Fig. S2f**). Instead the axonal arborization was significantly increased 100-200 μm from the soma (**Fig. 2g**), affecting tertiary and more distal axon branches (**Fig. S2j-m**).

**Figure 2.**
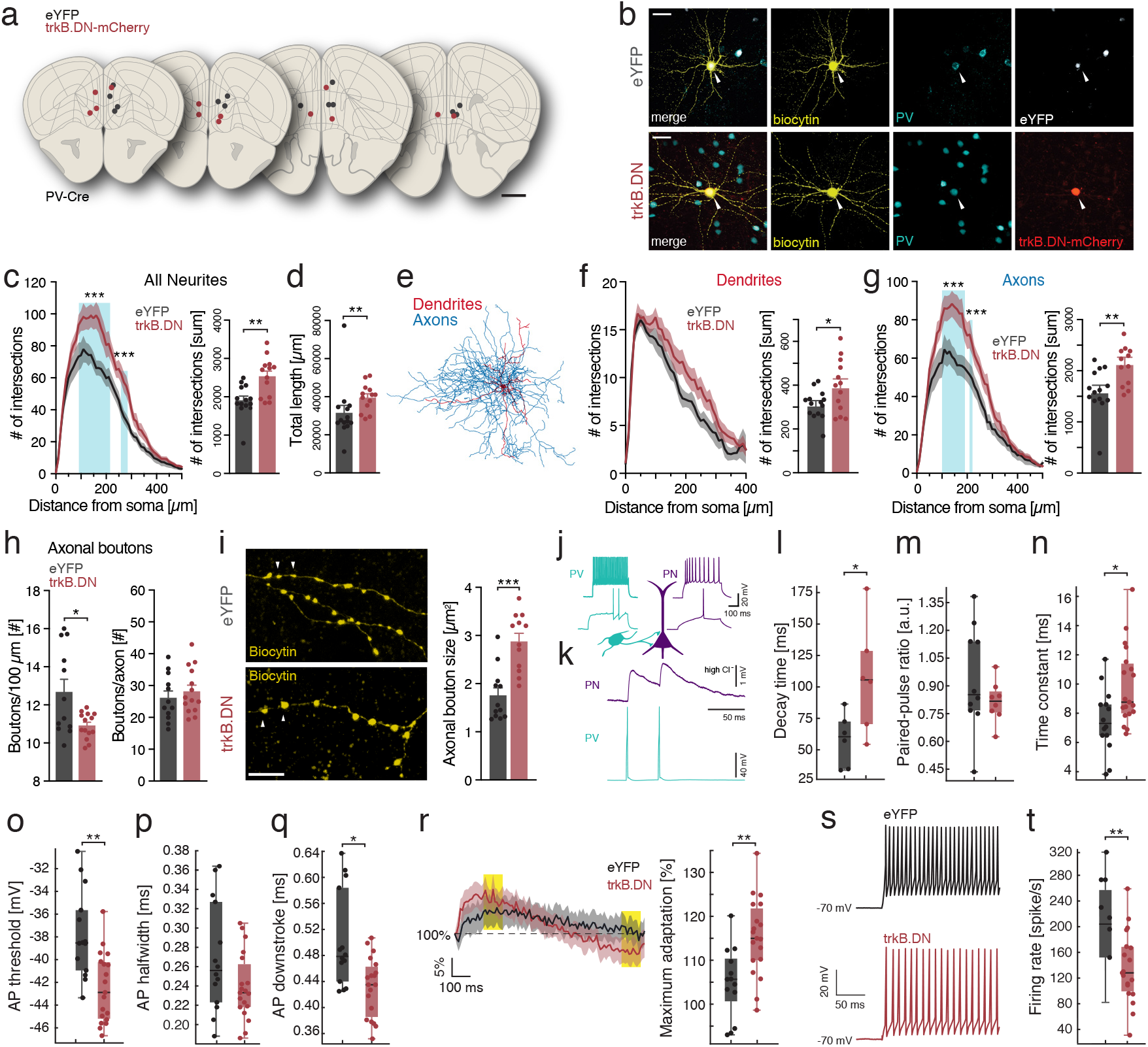
Morphological and functional changes in mPFC PV interneurons overexpressing trkB.DN. (**a**) Schematic coronal sections (2.3 to 1.7 mm anterior-posterior (AP) relative to Bregma) depicting the anatomical location of biocytin-filled PV interneurons subjected to morphological analyses. Red dots: mPFC interneurons expressing trkB.DN-mCherry (n = 13, in 6 trkB.DN mice). Gray dots: mPFC PV basket neurons and chandeliers expressing eYFP (n = 14, in 8 eYFP mice). (**b**) Representative biocytin-filled mPFC PV basked neurons stained for biocytin (yellow), PV (cyan), and eYFP (white) or mCherry (red), in an eYFP mouse (top), or a trkB.DN mouse (bottom). (**c**-**g**) Data based on Sholl analysis of the biocytin-filled, mPFC PV basket neurons in (a). (**c**) The neurites (dendrites + axons) of PV basket neurons expressing trkB.DN (red) display increased complexity compared to PV basket neurons expressing eYFP (gray), with a significantly increased number of intersections particularly 100-200 μm from the soma, and a significantly increased total number of intersections. eYFP: 1908 ± 109.9 intersections; trkB.DN: 2536 ± 140.8 intersections, *p* = 0.001. Blue shading: significant difference between trkB.DN and eYFP mice. (**d**) PV basket neurons expressing trkB.DN (red) have significantly increased total neurite length compared to PV basket neurons expressing eYFP (gray). eYFP: 31554 ± 3922 μm; trkB.DN: 39614 ± 2002 μm, *p* = 0.0013. (**e**) PV basket neuron with color-coding of the traced dendrites (red) and axons (blue). (**f**) The dendrites of PV basket neurons expressing trkB.DN (red) display increased complexity compared to PV basket neurons expressing eYFP (gray), as observed by an increased number of intersections at most distances from the soma, and a significantly increased total number of intersections. eYFP: 312.1 ± 17.15 intersections; trkB.DN: 395.8 ± 33.4 intersections, *p* = 0.0316. (**g**) The axons of PV basket neurons expressing trkB.DN (red) display increased complexity compared to PV basket neurons expressing eYFP (gray), with a significantly increased number of intersections particularly 100-200 μm from the soma, and a significantly increased total number of intersections. eYFP: 1601 ± 118.8 intersections; trkB.DN: 2153 ± 114.2 intersections. *p* = 0.0013. Blue shading: significant difference between trkB.DN and eYFP mice. (**h**) Quantification of axonal boutons in biocytin-filled mPFC PV basket neurons expressing trkB.DN (n = 14 neurons in 6 trkB.DN mice) or eYFP (n = 12 neurons in 8 eYFP mice). Left: boutons/100 μm, eYFP mice: 12.69 ± 0.66 boutons/100 μm; trkB.DN mice: 10.92 ± 0.18 boutons/100 μm; *p* = 0.0104. Right: the total number of boutons per axonal path, eYFP mice: 26.19 ± 2.1 boutons/axon, trkB.DN mice: 28.26 ± 1.93 boutons/axon; *p* = 0.4752. (**i**) Representative images of axonal boutons (arrow heads) in biocytin-filled mPFC PV basket neurons expressing eYFP (top) or trkB.DN (bottom). Expression of trkB.DN result in increased mean bouton size (eYFP mice: 1.76 ± 0.16 μm_2_, n = 12 axonal paths from 4 mice; trkB.DN mice: 2.88 ± 0.17 μm_2_, n = 12 axonal paths from 4 mice; *p* < 0,0001). (**j**-**t**) *Ex vivo* slice electrophysiology. (**j-m**) Paired recordings of PV interneurons and excitatory pyramidal neurons (**j**) Typical firing patterns at rheobase (bottom) and twice the rheobase current (top) of PV interneurons (green) and excitatory pyramidal neurons (PN; dark purple) in an eYFP mouse. (**k**) Representative traces of evoked APs in a mPFC PV interneuron (green) in a eYFP mouse, and the corresponding postsynaptic putative pyramidal inhibitory postsynaptic potential (IPSP) (dark purple). (**I**) Boxplots showing the decay time of the first IPSP in putative pyramidal neurons (eYFP mice (gray): 60.36 ms, n = 6 PV-PN pairs in 5 mice. trkB.DN mice (red): 105.58 ms, n = 6 PV-PN pairs in 6 mice), *p* = 0.0329. (**m**) Boxplots showing the paired pulse ratio of pyramidal neurons contacted by eYFP positive (grey; n = 10 PV-PN pairs in 8 mice) or trkB.DN-mCherry positive (red; n = 8 PV-PN pairs in 7 mice) PV interneurons. eYFP mice: 0.78; trkB.DN mice: 0.81; *p* = 0.6312. (**n**-**t**) Boxplots showing quantification of intrinsic properties of mPFC PV interneurons expressing YFP (n = 14 neurons, 10 mice), or trkB.DN (n = 19 neurons, 6 mice). (**n**) Membrane time constant. eYFP mice: 7.30 ms; trkB.DN mice: 8.76 ms; *p* = 0.01. (**o**) AP threshold. eYFP mice: −38.53 mV; trkB.DN mice: −42.88 mV; *p* = 0.0005. (**p**) AP width at half maximum. eYFP mice: 0.2560 ms; trkB.DN mice: 0.2330 ms; *p* = 0.07. (**q**) AP down stroke duration. eYFP mice: 0.4785 ms; trkB.DN mice: 0.4350 ms; *p* = 0.01. (**r**) Left: spike rate adaptation changes over 1 second current pulse at twice the rheobase current (grey: eYFP mice, n = 13 neurons, red: trkB.DN mice, n = 19 neurons). Right: boxplots of the maximum adaptation: change in firing rate between the immediate frequency around 200 ms and 930 ms (applied time points marked by yellow shading in left). eYFP mice: 105,0 %; trkB.DN mice: 115,3 %; *p* = 0.002 (**s**) Representative traces of action potentials (APs) at twice the rheobase current of mPFC PV interneurons in trkB.DN mice (red), and eYFP mice (gray). (**t**) Firing rate at twice rheobase current injection (eYFP mice: 204; trkB.DN mice: 128 spikes/s; *p* = 0.004). (d-h) Data shown as mean ± SEM. For boxplots (l-t), data shown as median, box: 25th and 75th percentile and whiskers: data points that are not outliers. For (r), data shown as mean and confidence interval. Unpaired t test, Two-tailed was used to assess significance if data passed the D’Agostino & Pearson normality test, if not, the Wilcoxon rank-sum test was used. For sholl analysis plot (c, f and g), significance was tested with Multiple t tests with the Holm-Sidak method. Scale bars: 1 mm in (a), 50 μm in (c). 10 μm in (i). * *p* < 0.05, ** *p* < 0.01, *** *p* < 0.001

TrkB signaling has been shown to be involved in the clustering and localization of GABAergic synapses (Xenos et al. 2018). We therefore next analyzed the inhibitory synapses by counting of axonal boutons, (indicative of synaptic terminals) of biocytin-filled PV interneurons, which revealed a significantly decreased density of boutons along the axons of trkB.DN expressing PV interneurons (**Fig. 2h**). However, accounting for their increased axonal length revealed that the total number of boutons did not differ between trkB.DN and eYFP mice (**Fig. 2h**), suggesting the presence of compensatory adaptations. Regardless, the boutons of trkB.DN expressing PV interneurons were rounded and corpulent, with significantly increased size, compared to the boutons of eYFP expressing PV interneurons (**Fig. 2i**).

### Changes in intrinsic properties and synaptic connectivity

We next used *ex vivo* patch-clamp recordings to investigate the effects of reduced trkB signaling on the synaptic connectivity between PV interneurons and target pyramidal neurons (PN) (n = 10 PV-PN pairs in 8 eYFP mice, n = 8 PV-PN pairs in 7 trkB.DN mice). Paired whole-cell recordings were used to probe the functional kinetics of synapses of mPFC PV interneurons terminating on pyramidal neurons (**Fig. 2j**). Action potentials (APs) were evoked in the PV interneurons and inhibitory postsynaptic potentials (IPSPs) were monitored in the postsynaptic PNs **(Fig. 2k)**. This identified that reduced trkB signaling in PV interneurons results in a significantly prolonged decay time of the IPSPs (**Fig. 2l**), suggestive of a deficiency in the regulation of the GABA release **(Fig. 2i)**. We used paired-pulse ratio (PPR) measurements to probe how reduced trkB signaling in mPFC PV interneurons affect short-term synaptic plasticity of PV-PN connections and found no difference in the PPR of PV-PN pairs between trkB.DN and eYFP mice **(Fig. 2m)**. Together the results indicate that reduced trkB signaling in mPFC PV interneurons results in prolonged PV-PN synaptic responses, but does not influence the amplitude of the responses, pointing towards primarily temporal imprecisions in the PV mediated inhibition of pyramidal neurons.

We thereafter investigated the intrinsic properties of mPFC PV interneurons (n = 14 in 10 eYFP mice, n = 19 in 6 trkB.DN mice). Characterization of the intrinsic excitability and firing properties revealed a significantly increased membrane time constant in PV interneurons expressing trkB.DN **(Fig. 2n),** indicative of increased response latency. We found the membrane capacitance in trkB.DN expressing PV interneurons to be significantly increased (**Fig. S2n**), whereas the input resistance did not change (**Fig. S2o**), attributing the altered membrane charging properties to increased capacitance. We measured a significantly lower rheobasic AP threshold in trkB.DN mice **(Fig. 2o)**, with a trend towards a narrower AP half-width **(Fig. 2p)** due to significantly faster AP downstrokes **(Fig. 2q)**. This threshold firing level data suggests increased excitability of mPFC PV interneurons expressing trkB.DN. However, probing of excitability on higher gain levels (by injection of larger depolarizing currents) showed a significantly higher maximum adaptation **(Fig. 2r)**, and significantly lower firing rate **(Figs. 2s, t)** in PV interneurons expressing trkB.DN. Thus, when under increased drive, trkB.DN-expressing PV interneurons maintained the firing rate less efficiently than PV interneurons expressing eYFP, possibly decreasing their inhibitory drive onto local excitatory neurons.

### Expression of trkB.DN in mPFC PV interneurons enhances aggression

Prior work has linked prefrontal E/I ratio alterations characterized by neuronal hyperactivity and decreased PV interneuron activity to abnormalities in social interaction (Yizhar et al. 2011; Ferguson and Gao 2018; Levy et al. 2019). Further, prefrontal reduction of the BDNF/trkB signaling has been associated with deficits in social interactions and increased aggression (Mikics et al. 2018). In line with these findings we observed that trkB.DN mice displayed increased aggression in social contexts (data not shown) and we, therefore, investigated social interaction directly by use of a well-validated test, the resident-intruder procedure (**Fig. 3a**, and **Methods**; Winslow, 2003). For scoring of baseline aggression, single adult PV-Cre mice (n = 19) were exposed to an unfamiliar juvenile intruder mouse (~4 weeks old) for four minutes, a social stimulus that normally does not elicit aggressive responses (Winslow 2003). Mice showing high levels of aggression were excluded (n = 2). The remaining mice were separated into two groups and injected with AAV-DIO-eYFP (n = 9 mice) or AAV-DIO-trkB.DN-mCherry (n = 8 mice) into the mPFC. All mice were thereafter subjected to the resident-intruder procedure on four additional occasions over the next 49 days, using different intruders at every occasion (**Figs. 3a**, **S3a**). Social behaviors (sniffing, tail rattling, aggression) and non-social behaviors (cage exploration, digging and grooming) were scored for all five test occasions, with a focus on the longest time point after viral injections (day 49 (d49); post; **Fig. 3a**). No significant behavioral differences were detected between the two groups of animals, except for measures of aggression (**Figs. 3b-g**, **S3b-h**). TrkB.DN mice spent more time engaged in aggressive behavior compared to eYFP mice at all tested time points (**Fig. S3b**), with a significant increase d49 (**Figs. 3c, d**). TrkB.DN mice also conducted a higher number of aggressive bouts (**Figs. 3e, f**), with a shorter latency to the first attack, compared to eYFP mice (**Fig. 3g**). TrkB signaling mediates effects of stress (Notaras and van den Buuse 2020), notably in the mPFC (Barfield and Gourley 2018). The same mice were therefore subjected to the open field test and tested in the elevated plus maze (**Fig. S3a**). No differences in locomotion were observed between trkB.DN and eYFP mice (**Fig. S3g**), and trkB.DN mice visited the center of the open field the first three minutes of the test as the eYFP mice (**Figs. S3h, i**). However, trkB.DN mice spent significantly less time in the center (**Fig S3j**), possibly indicating changes in anxiety. In support of increased anxiety, trkB.DN mice visited the open arms in the elevated plus maze significantly fewer times than eYFP mice, and also performed fewer head-dips over the edges of the open areas, a sign of increased anxiety (Rodgers and Dalvi 1997) (**Figs. 3h, i**). However, the total time spent in the open arms was not significantly changed (**Fig. 3j**). In summary, our findings directly link BDNF/trkB signaling in mPFC PV interneurons to aggression and point to a possible role in anxiety-related behavior.

**Figure 3.**
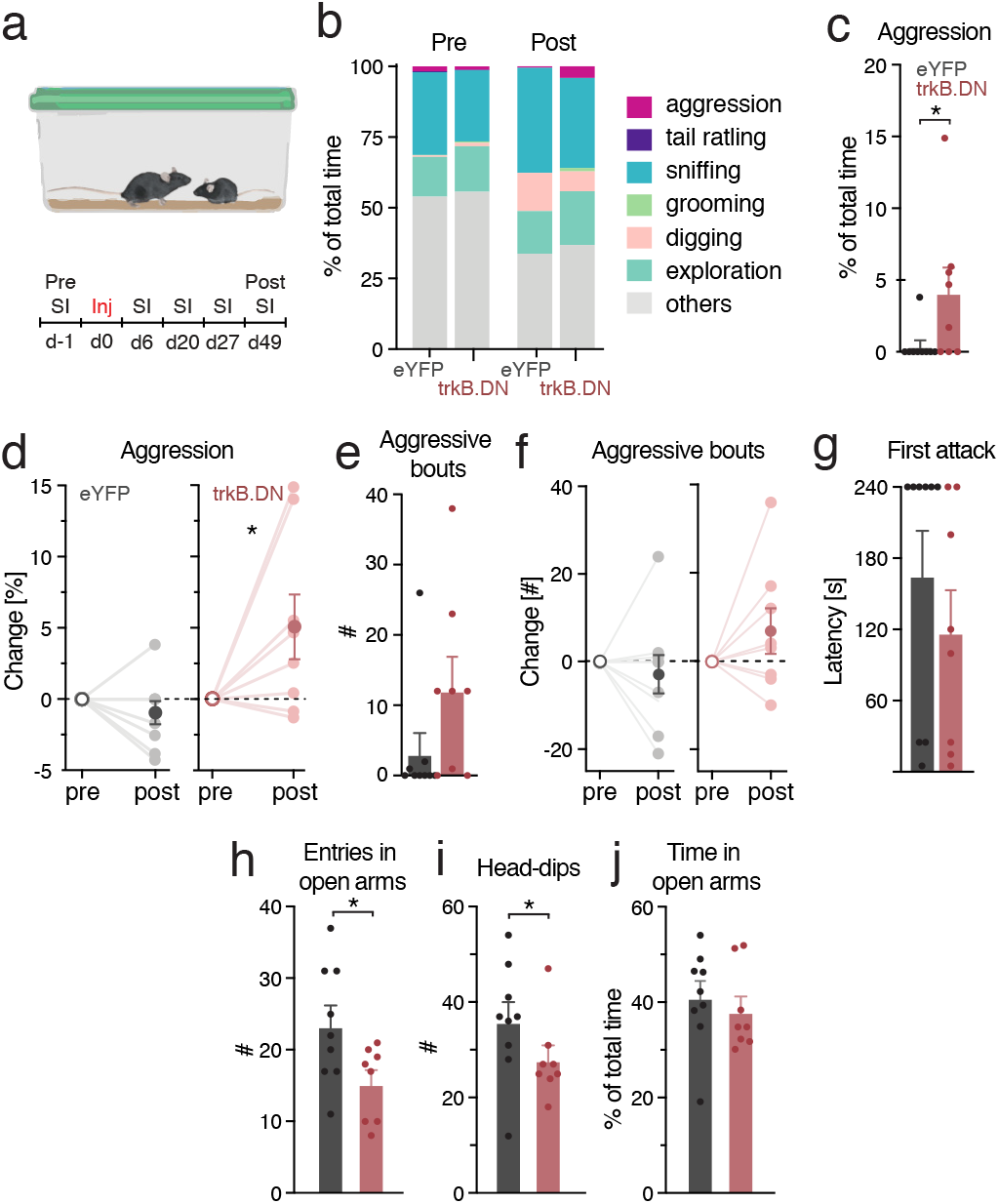
Expression of trkB.DN in mPFC PV interneurons enhances aggression. (**a-g**) Social interaction investigated by the resident intruder procedure. trkB.DN mice (n = 8 males), eYFP mice (n = 9 males). (**a**) The resident intruder procedure was conducted one day before viral injection and four times post injection. pre = before viral injection, post = 49 days after injection. (**b**) Percentage of total time spent in non-social (grooming, digging, cage exploration) and social behaviors (aggression, tail rattling, sniffing) for trkB.DN and eYFP mice during social interaction (4 min) pre (d1) and post (d49) injection. No behavioral differences were observed between trkB.DN and eYFP mice) before the viral injections (d1, pre): aggression (eYFP: 1.70 ± 0.87%; trkB.DN: 1.22 ± 0.35%; *p* = 0.6342), tail rattling (eYFP: 0.33 ± 0.33%; trkB.DN: 0.16 ± 0.16%; *p* = 0.6608), sniffing (eYFP: 29.27 ± 5.64%; trkB.DN: 25.11 ± 4.71%; *p* = 0.5849), grooming (eYFP: 0.14 ± 0.10%; trkB.DN: 0.48 ± 0.32%; *p* = 0.3022), digging (eYFP: 0.47 ± 0.38%; trkB.DN: 1.17 ± 0.68%; *p* = 0.3707), exploration (eYFP: 14.09 ± 4.74%; trkB.DN: 16.17 ± 2.99%; *p* = 0.7236) and others (eYFP: 53.99 ± 6.91%; trkB.DN: 55.69 ± 6.06%; *p* = 0.8578). (**c-g**) Expression of trkB.DN in mPFC PV interneurons results in increased aggression. (**c**) Percent time in aggression d49 (post): eYFP mice: 0.4 ± 0.4%; trkB.DN mice: 4.1 ± 1.7%, *p* = 0.0226. (**d**) Change (percent of time) in aggression (pre to post): eYFP mice: −0.9 ± 0.8%, *p* = 0.1875; trkB.DN mice: 5 ± 2.2%, *p* = 0.0391. (**e**) Number of aggressive bouts d49 (post): eYFP mice: 3.2 ± 2.9 bouts; trkB.DN mice: 12.25 ± 4.6 bouts; *p* = 0.0771. (**f**) Change (number) of aggressive bouts (pre to post): eYFP mice: −3.0 ± 4.3 bouts, *p* = 0.2891; trkB.DN mice: 6.9 ± 5.2 bouts, *p* = 0.1523. (**g**) Latency to first attack d49 (post): eYFP mice: 166 ± 37 seconds; trkB.DN mice: 118 ± 35 seconds, *p* = 0.2594. (**h-j**) Elevated plus maze. Same mice as in (a-g), performed d71. (**h**) TrkB.DN mice enter the open arms significantly fewer times than eYFP mice: eYFP mice: 23.44 ± 2.77 entries; trkB.DN mice: 15.38 ± 1.83 entries, *p* = 0.0320. (**i**) TrkB.DN mice make significantly fewer head-dips towards the open-arms than eYFP mice: eYFP mice: 36 ± 4 head-dips; trkB.DN mice: 28 ± 3 head-dips, *p* = 0.0490. (**j**) No difference was found in the time spent in the open-arms between trkB.DN, and eYFP, mice: eYFP mice: 41.10 ± 3.37 %; trkB.DN mice: 38.19 ± 3.06 % time spent in the open arms, *p* = 0.5360. Data shown as mean ± SEM. For bar plots, Unpaired t test, Two-tailed was used to assess significance if data passed the D’Agostino & Pearson normality test, if not Wilcoxon rank-sum test was used. For paired data, a Wilcoxon matched pairs test was used. * *p* < 0.05.

### Expression of trkB.DN in mPFC PV interneurons leads to changed network responses

To directly study mPFC responses and circuit dynamics during social interactions we conducted local field potentials (LFP) and single-unit activity recordings during the resident-intruder procedure (trkB.DN; n = 5, eYFP; n = 4 mice). Five weeks after viral injections, single-unit activity and LFP oscillatory pattern were recorded in the mPFC (**Figs. 4a** and **S4a**), before (9 min; baseline), during (4 min; social interaction: SI) and after social interaction (10 min; Post SI; **Fig. 4b**). As in the earlier experiments (**Figs. 3** and **S3**), the mice exhibited both social and non-social behaviors confirming that the tetrode implant did not overly affect the behavior **(Figs. 4c)**. Further, akin to our earlier results (**Figs. 3c-f**), trkB.DN mice exhibited increased aggression with a higher number of aggressive bouts compared to eYFP mice (**Fig. 4d**). No differences in locomotion between trkB.DN and eYFP mice were detected **(Fig. S4b)**. Analysis of the LFP oscillatory patterns (**Figs. 4e, f)** revealed that social interaction evoked increased power over a broad frequency spectrum in eYFP mice, with a significant increase specifically in the beta (12-24 Hz) and gamma (30-95 Hz) bands, an increase that became even more prominent after the intruder was removed (post-SI; **Figs. 4e, g**). In contrast, social interaction did, at large, not change the LFP power in trkB.DN mice (**Figs. 4f, g**). The only power change detected in trkB.DN mice was an increase in gamma activity after social interaction (post SI; **Fig. 4g**). Importantly, direct comparison of LFP oscillations between trkB.DN and eYFP mice revealed significantly increased *baseline* power over a broad frequency spectrum in trkB.DN mice **(Fig. 4h)**. Compared to the eYFP mice the trkB.DN mice, thus, presented enhanced baseline broadband activity and deficient induction of synchronous activity in response to social stimuli, particularly in the gamma band (**Figs. 4g, h**).

**Figure 4.**
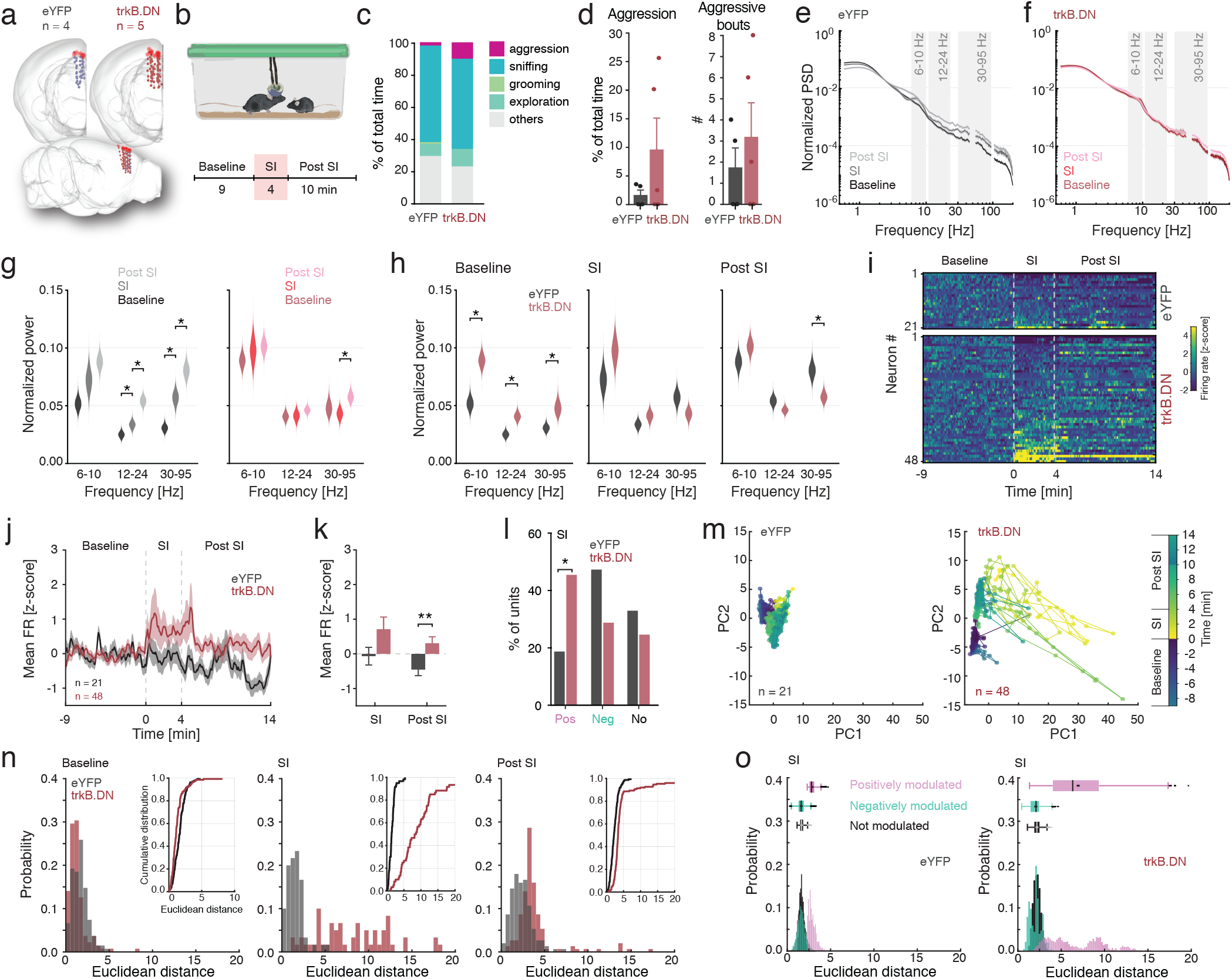
Expression of trkB.DN in mPFC PV interneurons leads to changed network responses during social interaction. (**a**) 3D illustrations of the reconstructed tetrodes positions. eYFP mice (gray; n = 4 males), trkB.DN mice (red; n = 5 males). Top: view from front, bottom: view from right side. (**b**) Top: illustration of implanted adult mouse, and juvenile intruder, respectively, in the resident intruder procedure. Bottom: Experimental timeline. Baseline: −9-0 min before introduction of the intruder; SI (social interactions) 0-4 min; Post SI (post social interaction) 4-14 min after removal of the intruder. The test was performed d50-60. (**c**) Percentage of total time spent in non-social (grooming, digging, cage exploration) and social behaviors (aggression, tail rattling, sniffing) for trkB.DN and eYFP mice, respectively, during the four minutes of social interaction. Due to lower n in both groups, no behavioral differences were detected between trkB.DN and eYFP mice. Sniffing: eYFP: 60.3 ± 6.4%; trkB.DN: 56.1 ± 8.2%, *p* = 0,7125; grooming: eYFP: 0.8 ± 0.5%; trkB.DN: 0.0%, *p* = 0,1279; exploration: eYFP: 7.6 ± 2.1%; trkB.DN: 11.0 ± 5.6%, *p* = 0,6242; others: eYFP: 29.6 ± 4.7%; trkB.DN: 23.1± 5.1%, *p* = 0,3907. No digging or tail rattling was observed. (**d**) Left: percentage time in aggression: eYFP mice: 1.7 ± 1.0%; trkB.DN mice: 9.8 ± 5.5% of time, *p* = 0,2452. Right: number of aggressive bouts: eYFP mice: 1.8 ± 0.5; trkB.DN mice: 3.2 ± 0.7 bouts; *p* = 0.503. (**e-o**) Chronic extracellular single-unit and LFP recordings during social interaction. (**e-f**) Bootstrapped normalized power-spectral-density (PSD) of the LFP for baseline (−9-0 min), social interaction (SI; 0-4 min), and post social interaction (Post SI; 4-14 min) for eYFP (**e**) and trkB.DN mice (**f**). Gray shading: the theta (6-10 Hz), beta (12-24 Hz) and gamma band (30-95 Hz), respectively. (**e**) Social interaction increases the power across many frequencies in eYFP mice, with further increases post SI. (**f**) Social interaction induces limited power changes in trkB.DN mice. (**g-h**) The distribution of the PSD in specific frequency bands (theta: 6-10 Hz, beta: 12-24 Hz, gamma: 30-95 Hz) during baseline, SI, and Post SI for eYFP (grays) and trkB.DN (reds) mice. (**g**) Direct comparison of the PSDs in the three frequency bands between eYFP mice (gray) and trkB.DN mice (red) during baseline, SI, and Post SI. trkB.DN mice have significantly increased baseline activity in all three frequency bands compared to eYFP mice. Social interaction increases the power in the eYFP mice, but not in trkB.DN mice, diminishing the power difference between the two groups of mice. The power increases further in eYFP mice after social interaction, with eYFP mice displaying significantly higher gamma power than trkB.DN mice post SI (despite the baseline gamma power being significantly higher in trkB.DN mice than in eYFP mice). Individual data point and confidence interval values for theta, beta and gamma range power intergroup comparisons during baseline, SI or Post SI can be found in the Supplementary table 1. (**h**) Direct comparison of the PSDs in the three frequency bands during baseline, SI, and Post SI for eYFP mice (left), and trkB.DN mice (right), respectively. Social interaction significantly increases the power in the beta and gamma bands in the eYFP mice. The only significant change in the trkB.DN mice is an increase in gamma power after social interaction (Post SI). Individual data point and confidence interval values for theta, beta and gamma range power intragroup comparisons during baseline, SI or Post SI can be found in the Supplementary table 1. (**i-o**) Analyses of the WS neurons. (**i**) The firing rate (z-scored) of individual WS neurons in eYFP mice (top: n = 21 neurons) and trkB.DN mice (bottom: n = 48 neurons) during baseline, SI, and Post SI. Dashed lines outline the SI period. (**j-k**) Mean firing rate (z-scored) of the WS population in the eYFP and trkB.DN mice, respectively. (**j**) Social interaction increases the mean firing rate of WS neurons in trkB.DN mice, but not in eYFP mice: eYFP mice: −0.07 ± 0.25, trkB.DN mice: 0.71 ± 0.35, *p* = 0.08 (baseline compared to SI). (**k**) After social interaction (Post SI) the mean WS firing rate is significantly higher in trkB.DN mice than in eYFP mice: eYFP mice: −0.45 ± 0.17; trkB.DN mice: 0.31 ± 0.18, *p* = 0.001. (**I**) A significantly higher proportion of the WS neurons is positively modulated by social interaction in trkB.DN mice than in eYFP mice: eYFP mice: 19% (4 neurons); trkB.DN mice: 46%, (22 neurons), *p* = 0.03, **χ**2 = 4.46). A lower proportion (non-significant) of the WS neurons is negatively modulated by social interaction in trkB.DN mice than in eYFP mice: eYFP mice: 48% (10 neurons); trkB.DN mice: 29% (14 neurons), *p* = 0.14, **χ**2 = 2.19. Not modulated neurons: eYFP mice: 33% (7 neurons); trkB.DN mice: 25% (12 neurons), *p* = 0.48, **χ**2 = 0.51). (**m**) Two-dimensional projection of population dynamics trajectories for eYFP mice (left) and trkB.DN mice (right) along the timeline of the experiment (indicated by the colorbar). PC1 and PC2 explained, respectively, 23.4% and 14.0% of variance seen in the eYFP mice, while PC1 and PC2 explained, respectively, 42.8% and 11.3% of variance in the trkB.DN mice. In eYFP mice, trajectories formed by different experiment periods seem to overlap (right), while trkB.T1 mice trajectories presents a different pattern. (**n**) Comparison of the probability distribution of Euclidean distance for each cluster (baseline, SI and Post SI) during the social interaction experiment shows significant differences between groups, for baseline (left; inlet: CDF comparison: *p* = 2.7×10_-6_), for SI (middle; inlet: CDF comparison: *p* = 3.15×10_-20_), and for post SI period (right; inlet: CDF comparison: *p* = 1.8×10_-15_). (**o**) Average bootstrapped Euclidean distances during social interaction, juxtaposing the positively, negatively and not modulated neurons for the eYFP (left) and trkB.DN (right) mice. The positive modulated neurons in trkB.DN mice display increased variability during social interaction. This increase is not present in eYFP mice. Data shown as mean ± SEM for (d, j-k). Data shown as mean ± SD for (e-h). For (l), the Chi-squared proportion test was used. For cumulative distribution function (n), the Kolmolgorov-Smirnov test was used. For bar plots (d and k) Unpaired t test - two-tailed was used to assess significance if data passed the Kolmogorov-Smirnov normality test, if not the Wilcoxon test was used. For paired data in (k), the Wilcoxon signed rank test was used. For (e and h) confidence intervals of bootstrapped data were used to assess significance. Scale bar: 100 μm in (b). * *p* < 0.05, ** *p* < 0.01.

We next analyzed single-unit activities during social interaction. The recorded mPFC units were pooled and classified into narrow spiking (NS) putative inhibitory interneurons (n = 18) and wide spiking (WS) putative pyramidal neurons (n = 69) based on spike waveform features (Ardid et al. 2015 and **Methods**). The units were thereafter separated based on animal ID (trkB.DN (n = 48 WS, 13 NS) vs eYFP mice (n = 21 WS, 5 NS); **Figs. S4d-f**.) excluding the NS, putative PV, units from further analyses due to their low number. No significant differences were detected in the baseline activity (mean firing rate and the coefficient of variation (CV) of the inter-spike interval) of the mPFC WS neurons between trkB.DN and eYFP mice (**Figs. S4g-h**). However, social interaction significantly increased the mean firing rate of the WS population in the trkB.DN mice, but not in the eYFP mice (**Fig. 4i**). Subdivision of the WS population based on the activity pattern of individual WS neurons during social interaction revealed that a significantly higher proportion of the WS neurons were positively modulated by social interaction in the trkB.DN mice compared to in the eYFP mice (**Figs. 4l** and **S4i-k**). In addition, a lower proportion of the WS mPFC neurons in the trkB.DN mice were negatively modulated by social interaction than in the eYFP mice (**Fig. 4l**). Nevertheless, the mean firing rate of the WS subpopulations (positively, negatively, and not modulated, respectively) did not differ between trkB.DN and eYFP mice (**Figs. S4i, j**), attributing the observed higher mean firing rate of the WS neurons in the trkB.DN mice (**Figs. 4j, k**) to a higher proportion of WS neurons being activated by social interaction in trkB.DN than in eYFP mice.

To further investigate how social interaction is represented in the mPFC WS population and to potentially reveal altered population-level dynamics in trkB.DN mice, we characterized the activity dynamics of simultaneously recorded WS mPFC neurons using neural trajectory analysis (Hamm et al. 2017; Pessoa 2019; Levy et al. 2019). After projecting the firing rate of the WS population along time (4 s bins) onto the first two principal components, we compared the intra-cluster variability of the three clusters representing the task intervals (baseline, SI, and post SI), by calculation of the distribution of Euclidean distance from the cluster centroid (**Figs. S4l-m**, and **Methods**). The variability in the distribution of the activity patterns of the WS population was significantly higher in trkB.DN mice than in eYFP mice in all three task intervals, with a remarkably pronounced difference in the WS activities during social interaction (**Figs. 4m, n**). To better understand the WS response variability in trkB.DN mice, we performed a bootstrapped version of the principal component analysis (see **Methods**) for separate analysis of the dynamics of negatively, positively, and not modulated WS neurons, respectively. This revealed that the largest divergence between the trkB.DN and eYFP mice was found in the dynamics of the positive modulated units, which displayed a marked variability in the average Euclidean distance, reflecting dynamics not present in the positively modulated WS population in eYFP mice (**Fig. 4o**).

We questioned if the altered population-level dynamics observed in trkB.DN mice during social interaction were generalized and would manifest also during other functional states engaging separate mPFC circuitry. Tail pinch is known to induce a state of arousal, characterized by desynchronized LFP activity, reduced power of slow-oscillations (0.5-2.0 Hz), and increased single-unit activity in the mPFC (Mantz et al. 1988; Massi et al. 2012). We recorded mPFC single-unit activity and LFP oscillations during a single tail pinch (7 s) in urethane anesthetized mice (eYFP: n = 3; trkB.DN: n = 5; **Figs. 5a**, **b**). As reported, tail pinch reduced the power of low-frequency oscillations (0.5-2.0 Hz) and we also observed an increase in higher frequencies in both groups (**Fig. 5c, d**). As before, the recorded single-units were classified into NS or WS neurons (**Figs. S5a-c**), and the activity of units with > 0.1 spikes/s during baseline (−2 to 0 min before tail pinch) was analyzed further (**Fig 5e;** eYFP: n = 45 WS; trkB.DN: n = 86 WS), again excluding the NS, putative PV, units due to their low numbers. No significant differences were detected in the baseline activity (mean firing rate and the coefficient of variation (CV) of the inter-spike interval) of the mPFC WS neurons between trkB.DN and eYFP mice (**Figs. S5d, e**). The tail pinch evoked increased firing, particularly in the trkB.DN mice (**Fig. S5f**), resulting in a significantly increased mean firing rate of the WS neurons in trkB.DN mice during the first 2 minutes post tail pinch compared to baseline (**Fig. 5f**). We again used the activity pattern of individual WS neurons to subdivide the WS population into positively, negatively and not modulated subpopulations. Paralleling the results derived by analysis of mPFC dynamics during social interaction, a significantly higher proportion of the WS neurons were positively modulated by tail pinch in the trkB.DN mice compared to in the eYFP mice (**Fig. 5g**). Further, a significantly lower proportion of the WS neurons in trkB.DN was negatively modulated by the tail pinch (**Fig. 5g**). Again, as seen during social interaction, the mean firing rate of the subpopulations of WS neurons (positively, negatively, and not modulated, respectively) did not differ between the two animal groups (**Fig. S5g)**. We next performed trajectory analysis of the WS population activity, as before (**Figs. 4m-o, S4l, m**). The firing rate of the WS population was projected along time (1 s bins) onto the first two principal components (**Fig. 5h**), and the intra-cluster variability of the clusters representing three temporal intervals (−2-0 min (baseline), 0-2 min, and 2-4 min; start of tail pinch = 0 min) was compared. Our analyses revealed abnormalities in the population-level dynamics of mPFC WS neurons in trkB.DN mice similar to what seen during social interaction, with the tail pinch (0-2 min, and 2-4 min) significantly increasing the variability in the distribution of the activity patterns in trkB.DN mice compared to eYFP mice (**Figs. 5h, i** and **S5h**). The bootstrapped version of the principal component analysis revealed, again, that the largest divergence between the trkB.DN and eYFP mice was found in the dynamics of the positive modulated units, which displayed dynamics and variability in the average Euclidean distance not observed in the positively modulated WS population in eYFP mice (**Fig. 5j**).

**Figure 5.**
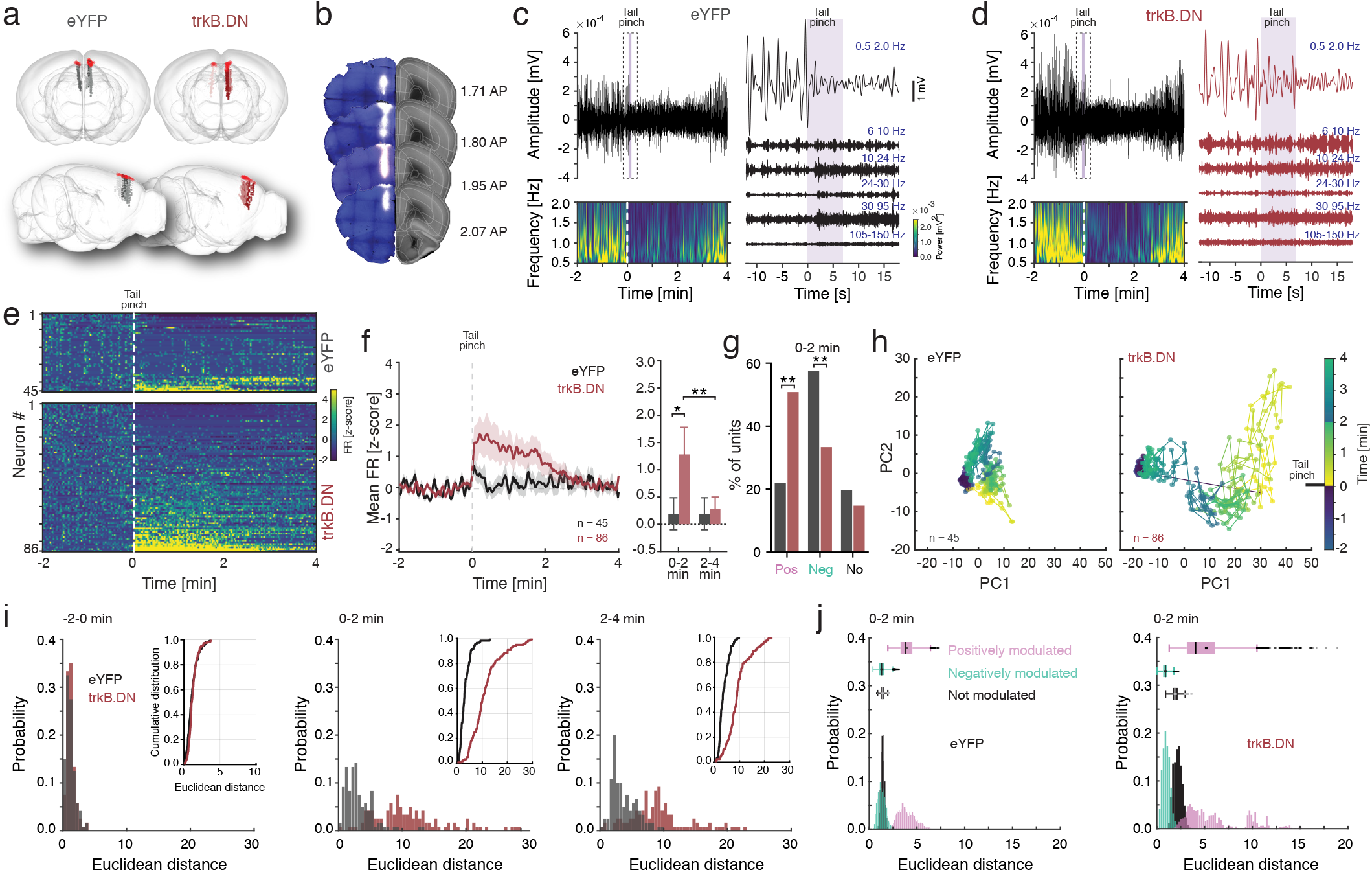
Expression of trkB.DN in mPFC PV interneurons leads to changed network responses during tail pinch. (**a-j**) Extracellular single-unit and LFP recordings under urethane anesthesia. eYFP mice (gray; n = 4 males), trkB.DN mice (red; n = 5 males). (**a**) 3D illustrations of the reconstructed probe positions. Top: view from front, bottom: view from right side. (**b**) Example brain sections from an eYFP mouse with DiI-labelling (white) of the tracts from the four-shank silicon probe. (**c, d**) Representative LFP recording from an eYFP (**c**), and a trkB.DN (**d**) mouse, respectively. Left: top; Filtered (0-200 Hz) LFP recording (6 min shown) before, during, and after tail pinch (7 s; violet bar). Dashed box: shown in closeup in right. Bottom: Wavelet spectrogram of representative LFP showing the decrease in power in slow frequencies (0.5-2.0 Hz) after tail pinch. Dashed line: start of tail pinch. Right: Close up of the dashed box in top left. Filtering of the LFP into different frequency bands. Violet bar: tail pinch (7 s). (**e**) The firing rate (z-scored) of individual WS neurons in eYFP mice (top: n = 45 neurons) and trkB.DN mice (bottom: n = 86 neurons) before, during and after tail pinch. Dashed line: start of tail pinch. (**f-g**) Mean firing rate (z-scored) of the WS population in eYFP and trkB.DN mice, respectively. (**f**) Left: Tail pinch significantly increases the firing rate of the WS population in trkB.DN mice (*p* = 0.0027) but not in eYFP (*p* = 0.0873, comparison −2-0 min vs 0-2 min). Dashed line: start of tail pinch. Right: The WS population in trkB.DN mice has a significantly higher firing rate than the WS population in eYFP mice 0-2 min after a tail pinch (eYFP mice: 0.19 ± 0.23; trkB.DN mice: 1.28 ± 0.46; *p* = 0.01), but not 2-4 min after the tail pinch (eYFP mice: 0.16 ± 0.20; trkB.DN mice: 0.28 ± 0.15; *p* = 0.51). The firing rate of the WS neurons in the trkB.DN mice decreases significantly after two minutes (*p* = 6×10_−5_) but not in the eYFP mice (*p* = 0.5918). (**g**) A significantly higher proportion of the WS neurons is positively modulated by social interaction in trkB.DN mice than in eYFP mice: eYFP mice: eYFP mice: 22% (10 neurons); trkB.DN mice: 51% (44 neurons), *p* = 0.001, **χ**2 = 10.21. A significantly lower proportion of the WS neurons is negatively modulated by social interaction in trkB.DN mice than in eYFP mice: eYFP mice: 58%, (26 neurons); trkB.DN mice: 34% (29 neurons), *p* = 0.008, **χ**2 = 7.01. Not modulated neurons: eYFP mice: 20% (9 neurons); trkB.DN mice: 15% (13 neurons), *p* = 0.48, **χ**2 = 0.50. (**h**) Two-dimensional projection of population dynamics trajectories for eYFP (left) and trkB.DN mice (right) along the timeline of the tail pinch experiment (timeline indicated by the colorbar). PC1 and PC2 explained, respectively, 25.2% and 19.2% of variance in the eYFP mice, while PC1 and PC2 explained, respectively, 68.4% and 8.6% of variance in the trkB.DN mice. (**i**) Comparison of the probability distribution of Euclidean distance for each cluster (−2 – 0 min, 0 – 2 min and 2 – 4 min) during the tail pinch experiment shows no difference during baseline (left; inlet: CDF comparison: *p* = 0.089), but trkB.DN Euclidean distance was significantly higher during the 0 - 2 min period after tail pinch (middle; inlet: CDF comparison: *p* = 1.97×10_-19_) and between 2 – 4 min period after tail pinch (right; inlet: CDF comparison: *p* = 2.13×10_-24_). (**j**) Average bootstrapped Euclidean distances during 0 – 2 min period after tail pinch, juxtaposing the positively, negatively and not modulated neurons for the eYFP (left) and trkB.DN (right) mice. The positive modulated neurons in trkB.DN mice display increased variability after a tail pinch. This increase is not present in eYFP mice. Data shown as mean ± SEM. For bar plots, the Wilcoxon rank-sum test was used to assess significance since data did not pass the Kolmogorov-Smirnov test for normality. For paired data in (f), the Wilcoxon signed rank test was used. For the Cumulative Distribution Function plots (i), Kolmogorov-Smirnov test was used to assess significance. * *p* < 0.05, ** *p* < 0.01.

## Discussion

Our results provide evidence for a critical role of BDNF/trkB signaling in the adult functions of prefrontal PV interneurons, with an impact on local network activities, including the E/I balance, population-level dynamics, and social behavior. Our experimental approach excludes alterations induced by changed differentiation or maturation of PV interneurons and associated secondary effects on the local mPFC network, and deficiencies caused by brain wide manipulations. Moreover, the approach enables observation of alterations induced by reduced BDNF/trkB signaling without deletions of the endogenous trkBs.

### Social interaction and sensory stimuli evoke altered network activity and excitatory responses

We found several alterations in the responses of mPFC excitatory WS neurons in trkB.DN mice during social interaction. While there was no difference in the baseline WS firing between trkB.DN and eYFP mice, social interaction significantly increased the mean firing rate of the WS population in trkB.DN mice, but not in eYFP mice. Moreover, the WS firing remained elevated for minutes. While the average firing rate of the WS populations positively, negatively, or not modulated by social interaction did not differ between trkB.DN and eYFP mice, a significantly higher proportion WS neurons was positively modulated by social interaction in trkB.DN mice than in eYFP mice, providing an explanation for the observed higher average firing rate of the total WS population in trkB.DN mice. Neural trajectory analysis through projection of the firing rate of the WS population as a function of time onto the two first principal components (PC) revealed that the population dynamics diverged between trkB.DN and eYFP mice. Particularly, in trkB.DN mice the trajectories during and after social interaction separated from the baseline trajectories, while in eYFP mice the three trajectories much overlapped. There was thus a higher variability in the WS population response to social interaction in trkB.DN mice than in eYFP mice. Interestingly, the increased response variability was specifically found in the positively modulated WS population in trkB.DN mice. Notably, very similar alterations in WS activity and dynamics in trkB.DN mice were seen in response to a tail pinch under anesthesia. While our experiments cannot reveal if individual neurons display similar responses to social interaction and tail pinch (as they were performed in different animal cohorts), the results indicate generalized alternations in the mPFC WS responses in disparate stimulus-driven brain states (social stimuli vs pure sensory stimuli - tail pinch), suggesting a general deficit in externally driven information processing (Hamm et al. 2017). Of note, all electrophysiological recordings and offline spike sorting were performed blindly with regard to the animal identity (trkB.DN vs eYFP), and disclosure of the identities revealed that a higher number of neurons were consistently recorded in the trkB.DN mice than in the eYFP mice, possibly due to the increased activity evoked in trkB.DN mice by social interaction and tail pinch allowing more neurons to be active enough to be recorded. We could not compare the activities of the putative inhibitory neurons (NS) between trkB.DN and eYFP mice due to the skewed neuron numbers which imposed restrictions on certain analyses.

Increased variability in stimulus responses in the visual cortex has been demonstrated in mouse models of schizophrenia when local multineuronal activity patterns were measured across time (using PCA-derived projection of the multineuronal state space onto three dimensions) (Hamm et al. 2017). Population states evoked by visual stimuli in wild type mice were unreliably activated in chronic mouse models of schizophrenia, suggesting a generalized disorganization of functional neocortical ensembles. Importantly, these alterations were not present in acute models of schizophrenia, including pharmacogenetic suppression of PV interneurons, indicating that specifically chronic models entail long-lasting structural changes in cortical networks that give rise to lasting disorganization of local neuronal ensembles (Hamm et al. 2017). Our long-term (several weeks) expression of trkB.DN in PV interneurons gives support for this notion. Recent work has directly investigated the role of PV interneurons in the regulation of cortical neuronal ensembles with potential relevance to our findings. First, optogenetic suppression of PV interneurons in the visual cortex during visual stimulation, but not during baseline, increased neuronal activity (Agetsuma et al. 2018). Similarly, we found no difference in the WS firing rate during baseline while both social interaction and tail pinch significantly increased the firing rate in trkB.DN but not eYFP mice. Further, in Agetsuma *et al*, PV suppression diminished the differential population activity patterns normally seen in response to different visual stimuli, i.e., the response reliability decreased, which reduced the functional repertoire of ensembles (Agetsuma et al. 2018). Reduced stimuli differentiation has also been demonstrated in the mPFC in a mouse model of social dysfunction associated with autism (Cntnap2 knockout) (Levy et al. 2019). As in our study the activity dynamics of simultaneously recorded mPFC neurons were characterized using neural trajectory analysis. Social and nonsocial stimuli (represented by olfactory cues) were found to evoke clearly divergent population responses in wildtype mice, a separation persisting seconds after stimulus offset. However, this category separation was decreased in the autism model, with mPFC neurons showing reduced differentiation between social and nonsocial stimuli (Levy et al. 2019). While we specifically assessed response variability, and not stimuli or category separation, the findings in the studies converge on disrupted population dynamics and disorganization of local population activity during stimuli processing. In both our study and in the study by Agetsuma *et al*, the altered dynamics were directly linked to changes in PV interneurons. Of interest, Agetsuma *et al* suggest that as the mechanisms underlying the alterations rely on anatomical network structure, the network aberrations are independent of the cognitive or behavioral state, and should present both in the awake state and during anesthesia. In support of this the activity deviations in trkB.DN mice manifested in both awake and anesthetized animals. Our *in vivo* recordings were performed during a single session of social interaction and did not allow analysis of mPFC population dynamics of specific behavioral variables during social interaction (aggression, sniffing, etc.). It would, however, be of interest to investigate if the population dynamics reflect the behavioral variables, and how reduced trkB signaling in PV interneurons affects stimuli separation.

Analysis of the LFP oscillatory patterns, which provides a proxy for the cortical activity levels and overall synchrony, revealed further alterations. The baseline LFP power in trkB.DN mice was significantly increased across a broad frequency spectrum compared to eYFP mice, including the theta, beta, and gamma bands. Interestingly, social interaction had little effect on the power spectra in trkB.DN mice while evoking a significant LFP power increase in eYFP mice, foremost in the beta and gamma bands. The beta and gamma power in eYFP mice were enhanced further after social interaction, resulting in a significantly higher gamma power than in trkB.DN mice. Thus, trkB.DN mice presented enhanced baseline broadband activity and deficient induction of synchronous activity in response to social stimuli, particularly in the gamma band. This deviates from what is observed in mice with global ablation of trkB in PV neurons during adolescence (crossing of PV-Cre mice to trkB_f/f_ mice), which leads to reduced cortical spontaneous LFP oscillations in frequencies >10 Hz in combination with reduced sensory-evoked gamma activity (Xenos et al. 2018). However, this model also entails severe structural and behavioral alterations, including loss of cortical volume with reduced number of PV interneurons and both motor and cognitive impairments. The gamma alterations (increased baseline and decreased induction) in the trkB.DN mice replicate findings in numerous studies of PV interneurons dysfunction, particularly NMDAR hypofunction models (Sohal et al. 2009; Korotkova et al. 2010; Yizhar et al. 2011; Carlén et al. 2012; Mathalon and Sohal 2015). PV interneurons’ decreased excitability and/or temporal dysfunctions of GABA release could be common denominators, as suggested by modeling (Jadi, Behrens, and Sejnowski 2016; Carlén et al. 2012; Bartos, Vida, and Jonas 2007 and see further below).

### trkB signaling and prefrontal PV interneurons in social behavior

The trkB.DN mice responded with increased aggression toward a juvenile intruder, and our PV specific genetic manipulation adds to a growing body of work coupling prefrontal BDNF/trkB signaling to social cognition and aggression. Highly aggressive rats have been demonstrated to display increased trkB.T levels and decreased trkB.FL/trkB.T ratio in the frontal cortex (Ilchibaeva et al. 2018). In line with this, increased aggression (induced by social isolation) in the resident intruder procedure has been linked to reduced levels of BDNF, increased levels of trkB.T1, and a decreased trkB.FL/trkB.T1 ratio in the infralimbic area of rats (Mikics et al. 2018). Mice with postnatal knockout of trkB in a majority of corticolimbic GABAergic interneurons display disrupted formation of social hierarchies, with elevated social dominance and aggression in male mice (Tan et al. 2018). Importantly, *in vitro* experiments demonstrated increased spontaneous excitatory synaptic transmission in the mPFC, and optogenetic silencing of the pyramidal neurons within the prelimbic cortex rescued the behavioral phenotype, coupling social behavior and trkB signaling to the E/I balance. Innovative optogenetic experiments in behaving mice have causally linked the E/I balance and social behavior (Yizhar et al. 2011). Optically enhanced excitability of excitatory (CaMKIIα-expressing) neurons, but not inhibitory PV interneurons, in the mPFC were shown to impair information transmission and social function. Of relevance to our findings, the increased E/I ratio was associated with enhanced mPFC firing and robust high-frequency power in the 30-80 Hz range. Further, the deficits in social behavior were ameliorated by optogenetic activation of mPFC PV interneurons (Yizhar et al. 2011). Our study provides *in vivo* data of altered prefrontal E/I balance and population dynamics underlying deficient social behavior, with a direct link to trkB signaling in PV interneurons.

### trkB signaling and intrinsic properties of prefrontal PV interneurons

While the regulatory role of BDNF/trkB signaling in the differentiation and maturation of PV interneurons has been firmly established (reviewed in Woo and Lu 2006), this role in the adult brain is unclear. Our experiments show that trkB signaling has a role in inhibitory synaptic modulation in the adult, and that reduced trkB signaling changes the intrinsic properties of PV interneurons. We find both pre- and postsynaptic alterations. Notably, several measurements indicated decreased responsiveness of PV interneurons expressing trkB.DN, including a significantly increased membrane time constant as a result of increased membrane capacitance. Further, under increased drive PV interneurons expressing trkB.DN displayed a significantly higher maximum adaptation and significantly lower firing rate, i.e., could not maintain the firing rate. However, the rheobasic AP threshold was significantly lowered, with significantly faster AP downstrokes, which would indicate increased excitability, possibly indicating compensatory adaptations. Investigation of postsynaptic excitatory neurons in the local network revealed primarily temporal imprecisions in the PV mediated inhibition in trkB.DN mice. The decay time of the IPSPs was significantly increased, indicating dysregulated GABA release in PV interneurons expressing trkB.DN, which could be a consequence of the discovered morphological malformation of the axon terminals. In coherence with our *in vivo* findings, the *ex vivo* data overall suggests improper activation of the PV interneurons and deficient PV mediated inhibition of postsynaptic excitatory neurons, which would be expected to enhance excitatory activities and negatively impinge on the E/I balance in the PFC. Interestingly, many of the alterations resemble what was found in the aforementioned genetic model with deletion of trkB in a majority of corticolimbic GABAergic interneurons, in which the trkB isoforms are deleted before the PV neurons have matured (Tan et al. 2018). The authors interpret the alterations as the PV interneurons being functionally immature. In a very recent study a novel optically activatable trkB was used for local and fast induction of trkB signaling in PV interneurons of the adult visual cortex (Winkel et al. 2020). 30-60 min after short-term optical trkB activation the excitability of PV interneurons was significantly decreased (through effects on Kv.3 channels). It is, thus, apparent that both increased and decreased trkB activity can affect the excitability of PV interneurons through multiple mechanisms, at different time scales. Compensatory adaptations are likely to be employed in response to long-term manipulations, but regardless, our data together with findings in genetic knockout models indicate that decreased responsiveness of PV interneurons is a lasting alteration. Our adult reduction of trkB signaling, however, rendered distinct morphological changes not seen in the genetic models. Postnatal knock-out of trkB in PV neurons decreases the number of neurites and the length and arborization of the neurites (Zheng et al. 2011; Tan et al. 2018; Xenos et al. 2018). We found increased neurite complexity, affecting both the dendrites and axons. The number, arborization, total length and the ending radius of the dendrites were significantly increased. The axons displayed a significantly increased complexity, with particularly increased arborization at distances from the soma where the complexity of PV interneurons overall is highest. This is also the location where PV basket neurons densely innervate local pyramidal neurons (Tremblay, Lee, and Rudy 2016). It is possible that the modifications found represent compensatory adaptations aimed to restore the decreased responsiveness and deficient regulation of excitatory postsynaptic responses. The viral construct employed in the current study can be used for further investigations of short-term or long-term alteration of trkB signaling in cell-types and brain regions of interest and should aid clarification of cell-autonomous trkB actions and compensatory adaptations. Further elucidation of how BDNF through interaction with a complement of receptor isoforms in different cell-types and subcellular compartments can instruct changes in morphologies and intrinsic properties is essential to further our understanding of how adult circuit functions are regulated by neurotrophic action.

## Materials and Methods

### Animals

All procedures were performed in accordance with the Guidelines of the Stockholm Municipal Committee for animal experiments (Stockholms djurförsöksetiska nämnd, approval N5/14). Animals used: Molecular and morphological investigations: adult (8-16 weeks old) male and female PV-Cre mice (n = 12; Jax Stock n° 008069). Ex vivo electrophysiology and morphology: adult (8-16 weeks old) male and female PV-Cre (n = 34), Lhx6-eGFP mice (n = 4; used to visualize Lhx6 expressing interneurons in control experiments; MMRC Stock n° 000246-MU) and Lhx6-eGFP/SST-Cre mice (n = 1; used to visualize Lhx6 expressing interneurons in control experiments; SST-Cre: Jax Stock n° 013044). In vivo electrophysiology and behavioral tests: adult (8-32 weeks old) PV-Cre male mice (n = 36). For the resident-intruder procedure, male juvenile (3-4 weeks old) PV-Cre mice were used as intruders. Single-nucleus RNA sequencing: adult (9 weeks-old) Vgat-Cre:H2B-eGFP mice (see further below). Mice were housed in groups (2–5 mice/cage) in individually-ventilated-cage systems (Sealsafe plus, GM 500, Tecniplast, Buggugiate, Italy), under standardized conditions with a 12-hour light-dark cycle (light 7:00 am), stable temperature (20 ± 1°C) and humidity (40-50%) with access to food (R70 Standard Diet, Lactamin AB, Vadstena, Sweden; 4.5 gm% fat, 14.5 gm% protein, 60 gm% carbohydrates) and water ad libitum. Implanted mice used for electrophysiology during behavior were single-housed after implantation (n = 10). All transgenic mice used were heterozygous for the transgene.

### Viral constructs and Cre-dependent gene expression

AAV-DIO-trkB.DN-mCherry, full name AAV-EF1a-DIO-trkB.DN-TM570-mCherry (Addgene, n°121502), was constructed by PCR amplification of rat truncated trkB.T1 with the sequence downstream of the transmembrane domain replaced by the sequence of three alanine residues followed by a stop codon (from the plasmid pLTM570, gift from Lino Tessarollo, NIH). For visualization of neurons expressing trkB.DN, the stop codon was removed and mCherry (from the plasmid pAAV-EF1a-double floxed-hChR2(H134R)-mCherry-WPRE-HGHpA, Addgene, no. 20297) was fused to the C terminal of trkB.DN, using the same open reading frame. trkB.DN-mCherry was inserted between the two double floxed sites in the pAAV-EF1a-DIO plasmid backbone (Addgene plasmid n°20949), using NheI and AscI as restriction enzyme cloning sites. AAV5 particles (8 × 10e12 viral particles/ml) were custom prepared by the Virus Vector Core Facility at the University of North Carolina at Chapel Hill. The AAV5-EF1a-DIO-eYFP (Addgene, n°27056) (3.3 × 10e12 particles/ml), was produced by the Virus Vector Core Facility at the University of North Carolina at Chapel Hill.

### Viral injections

A biosafety cabinet class 2 with sterile surgical environment was used for viral injections. Fixed to a Quintessential Stereotaxic Injector (Stoelting, Wood Dale, IL, USA), the micropipette (graduated borosilicate glass capillary; Wiretrol I, Drummond, Broomall, PA, USA) was filled with mineral oil, and 1 μL of virus mixed with Fast Green FCF Solution 2.5% (Electron Microscopy Sciences, Hatfield, PA, USA). The mouse was placed on a heating blanket (37 °C) in a stereotaxic apparatus (Harvard Apparatus, Holliston, MA, USA) and anesthetized (2% isoflurane). The mouse’s reflexes were tested and the isoflurane level was reduced to 1 or 1.5% over the course of the surgery. The scalp was shaved and lidocaine 1% was injected subcutaneously at the site of surgery. The mouse was treated with analgesic buprenorphine (0.2 mg/kg) before skin incision and the eyes were covered with eye lubricant (Viscotears, Novartis, France). Using a fresh scalpel blade, a single incision through the skin was made over the injection location. The connective tissue around the skull was gently removed for clear viewing of the Bregma. Bilateral craniotomies were performed over the mPFC (1.8 mm, ML: ±0.3 mm, DV: −2 mm relative to Bregma). Sterile saline 0.9% was regularly applied over the skull while drilling to remove bone dust and control heat generation. The total diameter of the burr hole was not larger than the diameter of the drill bit (around 100 μm). Using a syringe needle the remaining skull was removed, exposing the dura that was thereafter carefully opened. The tip of the micropipette was lowered through the dura to reach the target location (prelimbic region). 0.5 μL virus /hemisphere was injected at a rate of 0.1 μL/min. The micropipette was kept in place for 10 minutes after each finished injection and thereafter slowly retracted. The skin was closed with small amounts of cyanoacrylate glue (Vetbond Tissue Adhesive, Henry Schein, Melville, NY, USA), and carprofen (5 mg/kg IP; Norocarp, Norbrook Laboratories, Ireland) was given for analgesics before termination of anesthesia. The mouse was monitored until completely recovered and thereafter placed in the homecage. An additional dose of carprofen was administered 24 h after surgery.

### Tissue processing

The mice were deeply anesthetized with pentobarbital and transcardially perfused with 1x phosphate-buffered saline (PBS, pH 7.4) followed by 4% paraformaldehyde (PFA) in PBS (0,1M; pH 7.4). The perfused brain was removed from the skull and postfixed with 4% PFA in 0.1M PBS at 4°C for 16h. For vibratome cutting, the brain was thoroughly washed in 0.1M PBS and thereafter coronally sectioned (40 μm thickness) using a vibratome (Leica VT1200S, Leica Microsystems, Nussloch GmbH, Germany). The sections were stored in 1x PBS at 4°C. For Cryosectioning, brains were after post-fixing cryoprotected in 10% sucrose in 1x PBS overnight and thereafter frozen using liquid carbon dioxide. The brains were sectioned (14 μm thickness) using a CryoStarTM NX70 Cryostat (ThermoFisher Scientific, Waltham, MA, U.S.) and mounted on Superfrost Plus glass slides (ThermoFisher Scientific).

### Immunohistochemistry

#### General protocol

both vibratome cut (free-floating) and cryosectioned tissue were permeabilized for 1h with 0.3% TritonX-100 in 1x Tris-buffered saline (TBS, 38 mM Tris-HCl, 8 mM Trizma base, 120 mM NaCl in extra pure water), blocked with 10% normal donkey serum in 1x TBST for 1h at room temperature (RT), and thereafter incubated with primary antibodies (see Antibody list) in 1x TBST at RT for 12-24h. The sections were thereafter washed three times in 1x TBST, and incubated with a species-specific fluorophore-conjugated secondary antibody in 1x TBST for 3-5 h (see Antibody list for details). The sections were thereafter consecutively washed with 1x TBST, 1x TBS and 1x PBS (10 min each). Vibratome cut sections were mounted on glass slides (Superfrost Plus, ThermoFisher Scientific). All sections were cover-slipped (ThermoFisher Scientific) using 50:50 Glycerol:1x PBS. Images were acquired at 10 or 20x magnification with a fluorescent microscope (Leica DM6000B) connected to a Hamamatsu Orca-FLASH 4.0 C11440 digital camera (Hamamatsu, Hamamatsu City, Japan). The primary and secondary antibodies used are shown in the antibody list table.

#### Evaluation of cell-type specific expression of AAV-DIO-trkB.DN-mCherry

40 μm free-floating mPFC sections were stained with antibodies against PV and mCherry and mounted. Confocal images covering all six cortical layers were captured (LSM-800 Zeiss, Oberkochen, Germany) using a 20x or 60x objective and identical pinhole, gain, and laser setting. The co-expression of PV and mCherry was quantified using the Cell Counter plugin of the software FIJI (Schindelin et al. 2012).

#### GABA and PV levels and trkB EC/IC ratio

An antigen retrieval step was introduced before blocking during immunohistochemistry. For this the free-floating sections were incubated for 5 min in 70 °C warm sodium citrate buffer (10 mM tri-sodium citrate in distilled water, 0.05% Tween 20, pH 6), and thereafter washed in TBST (RT, 10 min). Immunostaining were thereafter performed as described above. The sections were mounted on glass slides (Superfrost Plus, ThermoFisher Scientific) using SlowFadeGold antifade reagent (ThermoFisher Scientific). Confocal z-stack images (6 images/stack) were captured (LSM-800 Zeiss) with a 60x objective and consistent settings. The corrected total cell fluorescence (CTCF) of GABA, PV, EC and IC was measured using the software FIJI. For antibodies, see Antibody list.

### Antibody list

#### Primary Antibodies

| Analysis | Primary antibody |  |  |  |  |
| --- | --- | --- | --- | --- | --- |
|  | Target | Host | Dilution | Company | Catalogue # |
| Virus specificity and efficiency | PV | guinea pig | 1:1000 | Synaptic Systems, Göttingen, Germany | 195 004 |
|  | mCherry | chicken | 1:500 | Novus Biologicals, Centennial, CO, USA | NBP2-25158 |
| Scholl analysis and axonal bouton counting | PV | guinea pig | 1:1000 | Synaptic Systems | 195 004 |
|  | mCherry | chicken | 1:1000 | Novus Biologicals | NBP2-25158 |
| GABA levels | PV | guinea pig | 1:1000 | Synaptic Systems | 195 004 |
|  | GABA | rabbit | 1:500 | Sigma-Aldrich, St. Louis, MO, USA | A 2052 |
|  | gephyrin | mouse | 1:500 | Synaptic Systems | 47 021 |
|  | mCherry | chicken | 1:500 | Novus Biologicals | NBP2-25158 |
|  | eGFP | chicken | 1:1000 | Abcam, Cambridge, United Kingdom | ab13970 |
| trkB.EC/IC ratio | PV | guinea pig | 1:1000 | Synaptic Systems | 195 004 |
|  | trkB-extracellular (AA 32-429) | rat | 1:300 | Antibodies-online, Aachen, Germany | ABIN1983534 |
|  | trkB-intracellular (AA 671-720) | rabbit | 1:300 | antibodies-online | ABIN1532221 |
|  | mCherry | chicken | 1:500 | Novus Biologicals | NBP2-25158 |
|  | eGFP | chicken | 1:1000 | Millipore, Burlington, MA, USA | ab13970 |

### Secondary Antibodies

| Analysis | Secondary antibody |  |  |  |  |  |
| --- | --- | --- | --- | --- | --- | --- |
|  | Target | Fluorophore | Host | Dilution | Company | Catalogue # |
| Virus specificity and efficiency | guinea pig | AlexaFluor488 | donkey | 1500 | Jackson ImmunoResearch, Cambridge, United Kingdom | 706-545-148 |
|  | rabbit | Cy3 |  | 1:500 |  | 711-175-152 |
|  | guinea pig | Cy3 |  | 1:1000 |  | 706-175-148 |
| Scholl analysis and axonal bouton counting | guinea pig | AlexaFLuor488 |  | 1:1000 |  | 706-545-148 |
|  | rabbit | Cy3 |  | 1:1000 |  | 711-175-152 |
|  | guinea pig | Cy3 |  | 1:1000 |  | 706-175-148 |
|  | Streptavidin, AlexaFluor 633 conjugate |  |  | 1:50000 | ThermoFisher Scientific | S21375 |
| GABA levels | guinea pig | DyLight™ 405 | donkey | 1:500 | Jackson ImmunoResearch | 706-475-148 |
|  | rabbit | Cy5 |  |  |  | 711-175-152 |
|  | mouse | AlexaFluor488 |  |  |  | 715-545-151 |
|  | mouse | Cy3 |  |  |  | 715-165-151 |
|  | chicken | Cy3 |  |  |  | 703-165-155 |
|  | chicken | DyLight™ 405 |  |  |  | 703-475-155 |
| trkB.EC/IC ratio | guinea pig | DyLight™ 405 |  |  |  | 706-475-148 |
|  | rat | AlexaFluor488 |  |  |  | 712-545-153 |
|  | rabbit | Cy5 |  |  |  | 711-175-152 |
|  | rabbit | Cy3 |  |  |  | 711-175-152 |
|  | rat | Cy3 |  |  |  | 712-165-153 |
|  | rabbit | AlexaFluor488 |  |  |  | 711-545-152 |

### *Ex vivo* electrophysiology

The animal was deeply anaesthetized and transcardially perfused with ice cold cutting solution, containing (in mM): 40 NaCl, 2.5 KCl, 1.25 NaH2PO4, 26 NaHCO, 20 glucose, 37.5 sucrose, 20 HEPES, 46.5 NMDG, 46.5 HCl, 1 L-ascorbic acid, 0.5 CaCl2, 5 MgCl2. The animal was then decapitated and the brain dissected out. 250 μm thick coronal slices were vibratome cut (VT1200S, Leica, Wetzlar, Germany) in ice cold cutting solution, the slices were incubated in cutting solution at 34°C for 13 minutes, and then maintained at room temperature in recording solution until recording, containing (in mM): 124 NaCl, 2.5 KCl, 1.25 NaH2PO4, 26 NaHCO3, 20 glucose, 2 CaCl2, 1 MgCl2.

For recording, the slices were superfused with extracellular solution kept at 33-35°C. Neurons were visualized using a 60x objective (Olympus, Tokyo, Japan) and a DIC microscope (Scientifica, Uckfield, UK), and fluorescent neurons identified using a pE-400 LED light source (CoolLED, Andover, UK). Patch pipettes (resistance 5-10 MΩ, pulled using a P-87 Flaming/Brown micropipette puller, Sutter Instruments, Novato, CA, USA) were filled with low chloride internal solution containing (in mM): 130 K-gluconate, 5 Na2-phosphocreatine, 1.5 MgCl_2_, 10 HEPES, 5 Mg-ATP, 0.35 Na-GTP, 1 EGTA, 8 biocytin. For part of the paired recordings a high-chloride internal solution was used, containing (in mM): 120 KCl, 5 Na_2_-phosphocreatine, 1.5 MgCl_2_, 10 HEPES, 5 Mg-ATP, 0.35 Na-GTP, 1 EGTA, 8 biocytin.

Signals were recorded with an Axon MultiClamp 700B amplifier and digitized at 20 kHz with an Axon Digidata 1550B digitizer (Molecular Devices, San Jose, CA, USA). Access resistance and pipette capacitance were compensated for. To assess passive and active membrane properties, neurons recorded in current-clamp mode were held at a membrane potential of −70 mV. The rheobase current was determined by applying near-threshold current steps, followed by steps proportional to the rheobase current with a duration of 1 s. An additional small hyperpolarizing step (30 pA) with a duration of 300 ms was used to determine the time constant and to monitor the access resistance throughout the recording. During paired recordings, a PV neuron (identified by red or green fluorescence and AP pattern) and a pyramidal neuron (identified by soma shape and AP pattern) were recorded in current clamp. A pair of APs were evoked in the PV neuron, with a 50-100 ms interval between the two APs and a 10 s interval between sweeps.

Intrinsic electrical properties, paired-pulse ratio, IPSP amplitude and IPSP slope were extracted using a custom-written Matlab (MathWorks, Natick, MA, USA) script, additionally using the Matlab scripts: boundedline (Copyright 2010, Kelly Kearney), plotSpread (Copyright 2017, Jonas), and normalitytest (Öner and Kocakoç 2017). The AP characteristics were determined from the first spike that occurred at rheobase current stimulation. Firing rate and adaptation were determined from a current injection of approximately twice the rheobase current. The amplitude of the first IPSP was defined as the difference between the baseline, measured over 15 ms (5 ms before the evoked spike), and the peak of the first IPSP. The slope of the first IPSP was fitted with an exponential function using physfit (Copyright 2006, Daniel Wagenaar). The amplitude of the second IPSP was determined by subtracting the fitted tail of the first IPSP from the peak of the second IPSP.

### Tissue clearing for Scholl analysis and counting of axonal boutons

Slices containing biocytin filled PV interneurons were fixed in 4% PFA (in 0.1 M PB, pH 7.8) overnight at 4 °C. The following day the slices were repeatedly washed in PB and thereafter incubated for 24 h in CUBIC reagent 1 (25 wt% urea, 25 wt% N,N,N’,N’-tetrakis(2-hydroxypropyl) ethylenediamine, 15 wt% polyethylene glycol mono-p-isooctylphenyl ether/Triton X-100). CUBIC is used to clear tissues by promoting the release of heme while maintaining a moderately alkaline environment (Susaki et al. 2014). The slices were thereafter washed three times in PBT (0.1 M PB containing 0.3 % Triton-X) and blocked for 3 h in 5 % donkey serum in PBT. The slices were incubated with primary antibodies in PBT (see Antibody list) for 48h at 4 °C. After washing with PBT the slices were incubated for 24 h at 4 °C in PBT containing 0.001 % secondary antibodies (see Antibody list) and 0.002 % streptavidin conjugated to Cy5 fluorophore (ThermoFisher Scientific). The slices were washed in PB for 3 h and thereafter incubated in CUBIC reagent 2 (50 wt% sucrose, 25 wt% urea, 10 wt% 2,2′,2′’-nitrilotriethanol, and 0.1% (v/v) Triton X-100) overnight and thereafter mounted on glass slides (Superfrost Plus, ThermoFisher Scientific).

### Scholl analysis and counting of synaptic terminals

Tiled confocal z-stack images (ca 60 images / stack; LSM-800, Zeiss, Oberkochen, Germany) were acquired of immunohistochemically stained biocytin-filled PV interneurons at 20x. The images were processed with Fiji software and the neurons reconstructed using the plugin Simple Neurite tracer, a semi-automated framework for tracing of neurons (Longair, Baker, and Armstrong 2011). The neurites were marked either as axons or dendrites and their length measured. Sholl analysis was used to characterize the neuronal arbors, in which a series of concentric (sampling) spheres (10 μm interval between the radii) were created around the middle point (the soma of the traced neuron) (Ferreira et al. 2014). The algorithm counts how many times neurites intersect the sampling spheres. In order to perform Sholl analysis on specific regions of the neurite arbor we used the conventional inside-out Sholl analysis where neurites that extend from the cell body are defined as primary neurites, neurites emanating from primary neurites defined as secondary neurites, and so on. Neurites classified as tertiary or higher were grouped together (O’Neill et al. 2015). Axonal boutons (putative synaptic terminals) were analyzed in the same images. For each neuron, 3 secondary axons with a length of 100-300 μm were randomly picked and all boutons were counted. The average number of boutons/100 μm was calculated for each axonal path. For calculation of the boutons’ size, the boutons were processed using the Pixel Classification workflow from the software ilastik (version 1.3.3) (Berg et al. 2019) and the results exported into the FIJI software where the average sizes of labeled pixels were calculated using the Analyze Particles plugin.

### Single-nucleus RNA sequencing

Vgat-Cre mice (Jax Stock No: 028862) were crossed to mice with Cre-dependent expression of histone-B associated eGFP (H2B-eGFP; RIKEN LARGE, CDB0203K) to generate mice with nuclear eGFP expression in Vgat-expressing neurons (Vgat-Cre:H2B-eGFP mice). The animals (n = 2 male, 9 weeks old) were killed with an overdose of isoflurane and the brains were rapidly extracted from the skull and immersed in an ice-cold aCSF solution. The brains were immediately sectioned (300 μm) in ice-cold aCSF using a vibratome (VT1200S, Leica), and the sections placed in a Petri dish with ice-cold Leibovitz’s L-15 medium (GIBCO, ThermoFisher Scientific). The PFC was dissected out from the sections and the tissue stored in 1 mL ice-cold Leibovitz’s L-15 medium containing 1 μL SUPERase RNase inhibitor (20 U/μl, ThermoFisher Scientific). Specifically, the neuron nuclei were isolated using a nuclear isolation protocol (Yeung et al. 2014). The tissue was homogenized in a 2 ml lysis buffer using a Dounce tissue grinder (7 ml, VWR). After addition of a 1.8 M sucrose solution (4 mL), the homogenized solution was added onto a sucrose cushion (2 ml) in a 10 ml Ultra-Clear centrifuge tube (Beckman Coulter). Separation of nuclei from the tissue was done by centrifugation (26.500xg at 4°C for 1.5 h) and after discarding of the supernatant the nuclei pellet was resuspended in a 500 μL Nuclear resuspension buffer. Single nuclei were isolated using Fluorescence-Activated Cell Sorting (FACS) by their GFP emission profile, and sorted into 384 well-plates containing 2.3 μL ice-cold lysis buffer. Plates containing nuclei were straightaway frozen on dry ice and stored on −80°C until additional processing. cDNA libraries were constructed and sequenced using a Smart-seq2 protocol (Picelli et al. 2013). Concisely, after lysis of the nuclei, the polyA RNA reverses were transcribed followed by PCR preamplification and purification. Tagmentation of the cDNA was followed by PCR amplification of the fragments and purification. The final cDNA libraries were sequenced using Illumina HiSeq 2000 (Illumina, San Diego, CA, USA). The reads were mapped and aligned to the mouse genome (mm10) and gene expression values were calculated as count values for each transcript. Analysis was performed on count values of the exome per nucleus. The sequencing data counts (from n = 648 individual nuclei) were analyzed using the Seurat package in R (versions 1.4.0 and 2.3.4, (Butler et al. 2018), as published before (Märtin et al. 2019). The count data were normalized (log2) and z-score of log(variance/mean) was used to identify variance genes. Cut off for high-low gene level was established at > 1 z-score. After performing a PCA to obtain the genes that are differentially expressed throughout the population, the significant Principal Components in the dataset were determined after random sampling with 1000 replicates and a projected PCA was used to expand the gene list and to avert losing possible marker genes. The resultant list of genes (3000 number of unique genes per nucleus from 648 Vgat-Cre:H2B-GFP nuclei) was once more analyzed for PCs and randomly sampled (1000 replicates). The resulting significant PCs were implemented into a non-linear dimensionality reduction (t-SNE) analysis. A subsequent density-based clustering was implemented and markers per cluster were identified based on their distinctive expression.

### Behavioral testing

All behavioral testing, and any included electrophysiology, and their analyses were performed by an experimented blind to the animal ID (trkB.DN or eYFP mice). All mice were habituated to the experimenter for 1–2 days prior to the first test day and transported to the testing room at least 1 h before any procedures to facilitate adaptation to the surroundings. All tests were conducted during the light phase, unless otherwise indicated.

### Resident-intruder test

The home cage (36 cm × 18 cm × 12 cm) of the resident mouse was used for the resident intruder procedure. The cage top was replaced by a transparent acrylic piece to allow video recording with a CCD camera. At the start of the session the food and enrichment were removed (original bedding remained) and the resident was free to explore the cage for 1 min. A juvenile (3-4 weeks old, group housed) male mouse was introduced in a corner opposite to the current resident’s location, kept in the cage for 4 min, and thereafter removed. Social and non-social behaviors (Winslow 2003) were scored post hoc from the videos using a custom-made Matlab (The MathWorks, Natick, MA, USA); exploration (searching environment or rearing along the cage border), digging (fast alternating movements of the forepaws scraping back material), grooming (mice in sitting position with licking of the fur, grooming with the forepaws, or scratching with any limb), tail rattling (fast waving movements of the mouse tail), sniffing (body trunk, anogenital and nosing exploration of the intruder) and aggressive (attack, fighting as kicking, biting, and wrestling) behaviors.

### Elevated plus maze test

Anxiety was probed in an elevated (73.5 cm above the floor) plus maze with four arms (each arm: length; 40 cm, width; 7 cm), with two of the arms holding 15 cm high opaque walls (closed arms). At the start of the test the mouse was placed in the center (5 × 5 cm) of the maze, facing one of the closed arms. The mouse was allowed for freely explore the maze for 5 min, and the behavior was recorded with a CCD camera. The behavior was analyzed with Biobserve Viewer Software (Biobserve GmbH, Germany) to score the number of entries and the time spent in the center, the closed arms, and the open arms, respectively. Head dips (downward movement of the head toward the floor in the room performed while in the open arms, a sign of reduced levels of anxiety) and stretches attended postures (stretching towards the open area from inside the closed area). The maze was cleaned with 70% ethanol and wiped dry with paper towels between each mouse.

### Open-field test

Locomotion and anxiety were scored in open field boxes (46 cm × 46 cm × 30 cm with 16 light beam arrays, ActiMot2, TSE Systems, Germany). The breaking of light beams was used to score the location and activity of the animal. In each session a single mouse was placed in the box (near a wall) and allowed to freely explore the box for 60 min. The total distance travelled was scored, as were the time spent, number of visits, and the velocity (cm/s) during visits, in the center of the box (imaginary square with 25% of the open-field area). The box was cleaned with 70% ethanol and wiped dry with paper towels between each mouse.

### Acute *in vivo* electrophysiology

The mouse was anesthetized with urethane (1.1 g/kg in sterile saline 0.9%, I.P.) 1 h prior to start of the recordings, fixed in the stereotaxic apparatus on a heating blanket with feedback loop able to maintain the body temperature at 37 ± 0.5 °C (Harvard Apparatus) and kept under isoflurane anesthesia (0.25%) during surgery. A craniotomy was made at the site of the drill hole made for viral injection, large enough to allow positioning of a four-shank silicon probe (A4×2-tet-5mm-150-200-312-A32, Neuronexus, Ann Arbor, MI, USA), which was coupled to an adapter for connecting a 32 channel headstage preamplifier (Neuralynx, Bozeman, MT, USA), linked to a Neuralynx Cheetah 64 system (Digital Lynx 4SX, Neuralynx). The dura was carefully removed. Small amounts of saline (0.9%) were continuously dropped on the craniotomy in order to prevent brain surface dryness. An additional craniotomy was made over the cerebellum for fixing of the ground-screw used as reference. Fluorescent lipophilic dyes (DiD or DiO, Vybrant™ DiO and DiD Cell-Labeling Solutions, ThermoFisher Scientific) were painted on the electrode shank for *post hoc* detection of the probe tracts. We let the electrode to air-dry for at least 5 min. The silicon probe was carefully positioned in the mPFC (AP: 1.75 mm, ML: ±0.3 mm, DV: −2 mm) and we monitored in real-time the electrophysiological signal for 30 min to allow the brain tissue to recover and optimize of the recording condition. Two different recordings were performed. First, one hour of recording was performed to monitor different oscillatory states induced by urethane (data not shown). Second, a 6 min recording was performed to access mPFC modulation induced by a tail pinch. After a 2 min baseline recording, a tail pinch (7 s) was manually applied using a plastic-coated surgical plier, and electrophysiological activity was recorded for the following 4 min. The recorded electrical signals were divided, pre-amplified (1000x) and filtered for LFP and single-unit activity. The biological signal was sampled at 32 kHz and band-pass filtered between 600–6,000 Hz to record spikes, and between 0.5–500 Hz to record LFPs. After the recordings the animal was deeply anesthetized with pentobarbital, transcardially perfused and the brain was recovered for *post hoc* detection of the probe location using SHARP-Track (Shamash et al. 2018) and SBA Composer (Bakker, Tiesinga, and Kötter 2015).

### Chronic *in vivo* electrophysiology

#### FlexDrive construction and implantation

For chronic *in vivo* electrophysiology flexDrive implants were constructed with 7 tetrodes (Voigts et al. 2013). Custom-made tetrodes consisted of four twisted fine wires (polyimide insulated ni-chrome wire, 12 μm, Sandvik-Kanthal, Stockholm, Sweden) that were gold-plated to reduce the impedance to 0.2-0.4 MΩ at 1 kHz. Seven movable tetrodes were loaded into medical-grade polyimide carrier tubes (0.005-inch OD, Phelps Dodge, Phoenix, AZ, USA) in the flexDrive. An extra wire was installed for electromyography (EMG) recording.

For implantation of the flexDrive, the mice were anaesthetized with isoflurane (2%) and the body temperature maintained at 37° with a temperature controller system. The animals were fixed in a stereotaxic frame and a hole was drilled through the skull (1.75 mm anterior to Bregma, and 0.30 mm lateral to midline). The flexDrive was positioned above the craniotomy and the tetrodes gradually lowered to the prelimbic area (1.25 mm ventral to the brain surface). Two miniature anchoring screws were used to attach the flexDrive to the skull (one on the anterior and one on the posterior part of the skull). Two Teflon coated stainless steel wires (0.005 inch bare, A-M Systems, Sequim, WA, USA) from the electrode interface board of the flexDrive were connected to the screws for grounding. The flexDrive was secured onto the skull using dental adhesive cement (Super Bond C&B, Sun Medical, Moriyama, Japan). The animals were injected with analgesic (Buprenorphine 0.1 mg/kg s.c.) at the beginning of surgery and carprofen (5 mg/kg; Norocarp, Norbrook Laboratories, Ireland) was intraperitoneally injected at the end. The animal was monitored until completely recovered from the anesthesia and thereafter single-housed. An additional dose of carprofen was administered 24 h after surgery.

### Electrophysiological recordings during the resident intruder procedure

The resident intruder procedure was performed as described above with the exception that the resident was connected to the Neuralynx Cheetah 64 system (Digital Lynx 4SX, Neuralynx) and allowed to explore the cage freely for 9 min before the intruder was introduced (baseline). The intruder was introduced (social interaction), and removed after 4 min. The recordings continued for 10 min after the intruder had been removed (Post SI). For synchronization to recorded neural data, a custom-made Python 3.7 (Python Software Foundation) script was used to control the CCD camera connected to a USB-controlled Arduino microprocessor (Arduino Uno Rev3, Arduino, Italy) that outputs a train of voltage pulses (TTL trigger directly to the neural acquisition system) during electrophysiology recording.

### Electrolytic lesions

Electrolytic lesions were performed after the chronic recordings for mapping of the tetrode recording sites. The mouse was anesthetized with isoflurane (3%) and a crocodile clamp was fixed on one of the ears of the animal and 30 mA current was applied between the crocodile clamp and target electrode channel on the tetrodes for 30 s per channel to produce an electrolytic lesion. The mouse was thereafter perfused and the brain recovered for *post hoc* detection of the location of the tetrodes using SHARP-Track (Shamash et al. 2018) and SBA Composer (Bakker, Tiesinga, and Kötter 2015).

### Data analysis

Data analysis was conducted using GraphPad Prism version 8.00 (GraphPad Software, La Jolla, CA, USA) or custom software written in Matlab (The MathWorks, Natick, MA, USA).

### Unit sorting and classification

Single-units were manually sorted and identified by various spike waveform features (energy, peak, area, valley, peakValleyRatio, spike width, peakIndex, peak6to11 and principal components) using the offline spike-sorting toolbox MClust version 4.4 (written by A. David Redish). Only well-isolated units (Schmitzer-Torbert et al. 2005) with isolation distance > 15, L-ratio < 0.2, and the spikes < 0.01% at ISI < 2 ms were included in the data analysis.

The unit classification was performed according to Ardid et al. 2015. First, we verified if peak-to-trough duration of all units followed a bimodal distribution through a Calibrated Hartigan’s’ dip test analysis. Second, two Gaussian Mixture Models (GMM) were fit to the peak-to-trough duration for objective classification in wide-spiking (WS) putative pyramidal neurons or narrow-spiking (NS) putative interneurons, resulting in a WS-likelihood and a NS-likelihood for all units. For WS classification, only units with a WS-likelihood 10 times larger than the NS-likelihood were selected. In a similar manner, for NS classification only units with a NS-likelihood 10 times larger than the WS-likelihood were selected. Units that did not meet this criterion were not included in analysis (and categorized as unclassified). The classification was performed with an Open Source toolbox available at https://bitbucket.org/sardid/waveformanalysis.

### Analysis of mPFC activities during social interaction

Recordings of LFP and single-unit activity were done during 3 separated periods: After a baseline period of 9 minutes, an intruder mouse was placed into the home cage and the social interaction lasted a total of 4 min. After the removal of the intruder, the recording continued for an extra period of 10 min.

For the analysis of the mPFC oscillations, we first calculated the LFP’s root mean squared (RMS) envelope and established a manual threshold for movement detection in order to avoid movement-induced noises. Then, the LFP signal was segmented into 10 seconds blocks, and blocks presenting values above the manual threshold were removed. We next calculated the mean normalized LFP power for different frequencies (Theta: 6-10 Hz, beta: 12-24 Hz and gamma range: 30-95 Hz) using a bootstrap method. For each iteration we randomly chose blocks (with reposition) from both eYFP and trkB.DN groups (n = 41 blocks for baseline, n = 21 for social interaction and n = 51 for post-social interaction) and calculated the normalized power. For each 10 seconds blocks, Power Spectral Density (PSD) was estimated using Welch’s method (Matlab algorithm), with 1 second Hamming window, with 50% overlap and a 5120 Fast-Fourier transform (FFT) points. Normalized power was obtained by dividing the PSD estimation by the integrated power over all frequencies (0-200 Hz). PSDs for random blocks were then averaged for each group at each iteration. The process was repeated 10000 times, and the confidence interval of PSD distribution at each frequency band was calculated. For comparisons (group or intragroup), difference of bootstrapped datasets and the respective confidence interval were calculated. Distributions where the confidence interval contained the value “zero” were not considered significantly different.

For single-unit activity analysis, firing rate histograms were calculated for each individual WS neuron during SI, with a 4 seconds bin size and convoluted with a Gaussian Kernel of 12 seconds. The firing rate was z-scored against the baseline period. Only WS neurons with a firing rate of at least 0.1 spike/s during baseline were selected to further analysis. The mean z-scored firing rate was compared between groups, as well as the mean firing rate (in spike/s) and the coefficient of variation (CV; defined as the standard deviation divided by the mean of inter-spike interval) of baseline activity.

Neurons were classified as negatively modulated, not modulated, and positively modulated by comparing all bins during the baseline with the first 2 minutes of social interaction with a Wilcoxon Rank Sum test. Neurons that presented a different distribution during social interaction in comparison with the baseline period (p < 0.05) and a higher mean firing rate were considered positive modulated neurons, while neurons that present a different distribution and a smaller average firing rate were considered as negative modulated. Neurons that did not present different distributions before and after the tail pinch were considered as not modulated. Proportion and mean z-scored firing rate of negatively modulated, not modulated, and positively modulated were compared between groups.

Mice movements during social interaction were analyzed using Deeplab Cut (Mathis et al. 2018). Four points over the mouse head (including the flexDrive) were used for position tracking. Training was performed with 20 frames of each recording using a *ResNet 152* pretrained network, performing 900000 iterations. After tracking analysis, the average velocity (in pixel/s) during baseline, social interaction and post social interaction, was inferred from average position between all markers.

### Analysis of mPFC activities during tail pinch

The mean firing rate was calculated for each individual WS neuron at three different intervals: −2 min to tail pinch (baseline; −2-0 min), from tail pinch to 2 min after start of tail pinch (0-2 min), and from 2 min to 4 min after tail pinch (2-4 min). Spike trains were discretized in 1 second bins, convolved with a Gaussian Kernel of 4 seconds. Firing rate was thereafter z-scored in relation to the baseline. Only neurons with mean firing rate higher than 0.1 spike/s during baseline were included in further analysis. All further analysis were conducted as for mPFC activities during social interaction,

### Neural trajectory analysis

For analysis of population neural trajectories, the first two principal components of the z-scored firing rates of WS neurons were calculated with a PCA for each temporal bin (SI: bins of 4 s; tail pinch: bins of 1 s) (Hamm et al. 2017; Pessoa 2019; Levy et al. 2019). Projecting the first and second principal components in a two-dimensional space, we compared the variability of resulting clusters representing task intervals (social interaction: baseline, SI and post SI; tail pinch: −2-0 min, 0-2 min and 2-4 min after tail pinch) in terms of Euclidean Distance. The Euclidean distance was calculated as the pairwise distance values of PC1 and PC2 for each cluster centroid. Using a Cumulative Distribution Function, we compared the probability distribution of Euclidean Distance for each cluster between trkB.DN and eYFP mice with a Kolmogorov-Smirnov test for continuous distributions. To analyze the trajectories of specifically the negatively, positively and not modulated neurons for each task interval, a bootstrapped version of the trajectory analysis was performed with 10000 iterations. At each iteration, a random sample of neurons (with repetition) were selected, and the resulting first and second PCs of z-scored firing rate was calculated. Differently from the analysis performed with all WS neurons, here, the average Euclidean distance was calculated for each iteration, resulting in a distribution of average Euclidean distances for the negatively, positively and not modulated WS neurons.

### Statistics

A D’Agostino & Pearson test or Kolmogorov-Smirnov was used to assess the homogeneity of variance. If the data fulfilled the criteria for normal distribution, we analyzed the data using parametric statistics. If the data didn’t pass the normality test, we used the Wilcoxon rank-sum test or the Wilcoxon matched-pairs signed-rank test. For cumulative probability distribution, the Kolmogorov-Smirnov test was used. Only values of p < 0.05 were considered statistically significant. For bootstrapped analysis, only confidence intervals not containing zero were considered statistically significant.

## Author contributions

M.C and K.M conceptualized the study. Y.X. and K.M. designed and cloned the AAV-DIO-trkB.DN-mCherry. N.G. and J.I. designed and performed molecular experiments and analyzed data. J.A.v.L., J.F. and M.Z. designed and performed the ex vivo electrophysiology experiments and analyzed the data. N.G. and C.L.A. designed and performed the behavioral experiments and analyzed the data. N.G., L.R.Z. and C.L.A. designed and performed in vivo electrophysiology experiments and analyzed data. H.K. performed in vivo electrophysiology data analysis. A.M. performed the scRNAseq and analyzed the data. N.G., L.R.Z., J.A.L, A.M., C.L.A. and M.C. prepared figures. N.G., L.R.Z, C.L.A and M.C. wrote the original draft, N.G., L.R.Z. and M.C. wrote the final manuscript, and all authors reviewed the final manuscript.

## Acknowledgements

We would like to thank Lino Tessarollo and Mary Ellen Palko for generously donating the pLTM570 plasmid; Maria Lindskog, André Fisahn and Richard Andersson for advices on *ex vivo* electrophysiology experiments; Moritz Weglage for helping with the behavioral recording setup. Cantin Ortiz for helping with the social interaction analysis custom-made script. Image of neurons in Fig 1a was adapted from Federico Claudi, SciDraw (https://scidraw.io/).

---

**Figure S1.**
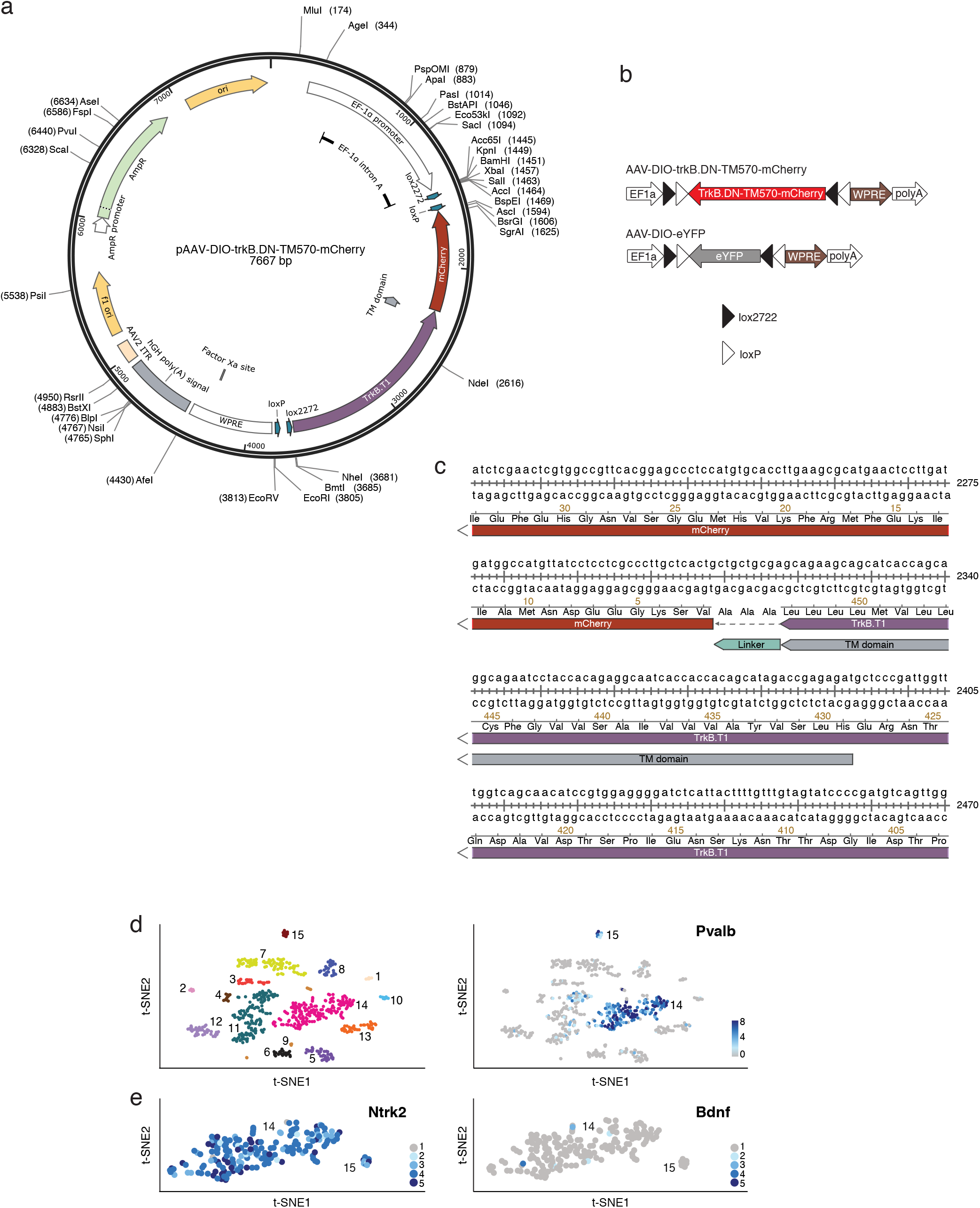
Virus information and trkB and BDNF levels in PV interneurons. (**a**) Full Sequence Map of the vector pAAV-DIO-trkB.DN-TM570-mCherry for Cre-inducible expression of trkB.DN-mCherry (Image created with SnapGene). (**b**) The viral vectors used in the present study. AAV-DIO-trkB.DN-TM570-mCherry (full name AAV5-Ef1a-DIO-TrkB.DN-TM570-mCherry): AAV vector with Cre-dependent expression of a truncated version of the trkB receptor fused to mCherry, AAV-DIO-eYFP (full name AAV5-Ef1a-DIO-eYFP): AAV vector with Cre-dependent expression of eYFP. L-ITR and R-ITR: Left and Right Inverted Terminal Repeat (ITR) sequences; EF1a: elongation factor 1a promoter; WPRE: Woodchuck hepatitis virus Posttranscriptional Regulatory Element. (**c**) Highlight of the sequence part around the link of mCherry and trkB.T1 after the transmembrane domain (TM domain) of trkB.T1, forming the vector AAV-DIO-trkB.DN-mCherry. (**d**) Left: visualization of the molecular diversity of mPFC GABAergic (Vgat-expressing) neurons in a t-SNE plot. Each dot represents an individual GABAergic neuron nucleus (n = 648 nuclei). Right: t-SNE plot of the distribution of the expression of *Pvalb* in the clusters in left. Two clusters, cluster 14 (n = 175 nuclei) and 15 (n = 12 nuclei) express high levels of *Pvalb* (normalized log2-expression). (**e**) Distribution of the expression of *Ntrk2* (left) and *Bdnf* (right) for the *Pvalb* expressing clusters 14 and 15 (normalized log2-expression). 98.9% (173/175) of the nuclei in clusters 14, and 100% (12/12) of the nuclei in cluster 15, express *Ntrk2*. In contrast, there is a very limited expression of *Bdnf* in the *Pvalb* expressing neurons; 2.9% (5/175) of the nuclei in cluster 14, and 0% (0/12) of the nuclei in cluster 15, express *Bdnf*.

**Figure S2.**
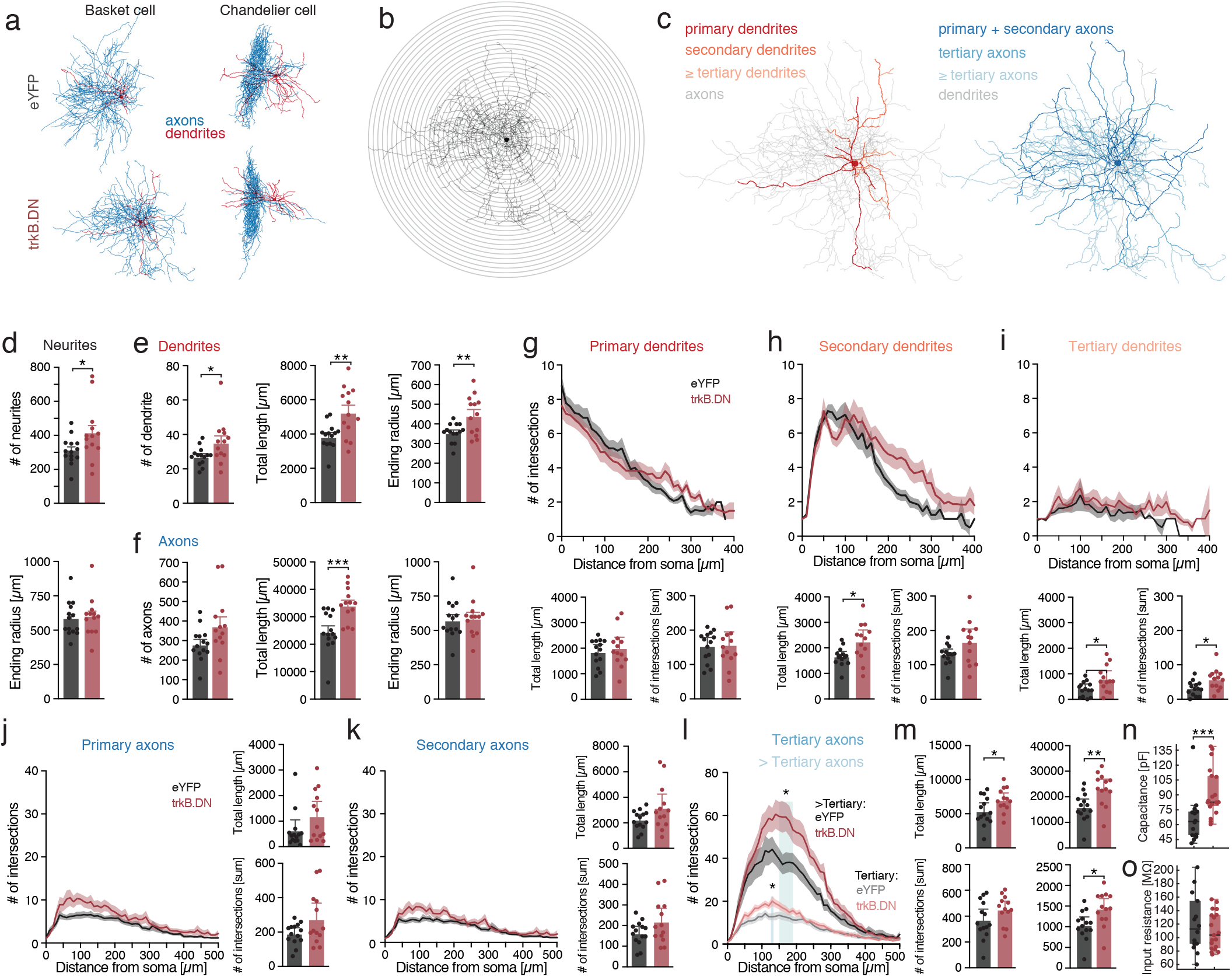
Morphology and *ex vivo* recordings. (**a**) Representative reconstructions of biocytin-filled mPFC PV basket cells and chandelier neurons, respectively, in eYFP mice (top) and trkB.DN mice (bottom). Blue: axons, red: dendrites. (**b**) Sholl analysis. Digitally applied concentric rings spaced 10 μm apart are centered on the soma center and the number of intersections between the neuronal process and ring at a given radius are counted. An estimate for morphological attributes of the neuron, as the number of processes, their length, but also the distance of the last ring intersected by a process (ending radius) are then analyzed. (**c**) Reconstructed biocytin-filled mPFC PV basket neurons with color coding of the dendritic (left) and axonal (right) hierarchical branching. Primary neurites (axons and dendrites): extending from the cell body; secondary neurites: emanating from primary neurites; tertiary neurites: emanating from secondary neurites, and so on. (**d**-**m**) Data based on Sholl analysis of the biocytin-filled, mPFC PV basket neurons in Fig. 4a. eYFP mice: n = 14 neurons in 8 mice; trkB.DN mice: n = 13 neurons in 6 mice. (**d**) Top: the number of neurites (eYFP mice: 310.9 ± 21.58 paths; trkB.DN mice: 413.5 ± 45.49 paths; *p* = 0.0474). Bottom: The ending radius of the neurites (eYFP mice: 582.1 ± 33.53 μm; trkB.DN mice: 598. 5 ± 40.19 μm; *p* = 0.7281) for all neurites combined. (**e**) The number of dendrites (eYFP mice: 27.43 ± 1.41 dendrites; trkB.DN mice: 35.69 ± 3.54 dendrites; *p* = 0.0236), total dendritic length (eYFP mice: 3896 ± 199.8 μm; trkB.DN mice: 5290 ± 405 μm; *p* = 0.0041) and ending radius of the dendrites (eYFP mice: 357.1 ± 12.69 μm; trkB.DN mice: 446.2 ± 27.65 μm; *p* = 0.0061). (**f**) The number of axons (eYFP mice: 283.5 ± 22.2 axons; trkB.DN mice: 377.8 ± 43.95 axons; *p* = 0.0614), total axonal length (eYFP mice: 24863 ± 1932 μm; trkB.DN mice: 34337 ± 1705 μm; *p* = 0.0002), and ending radius of the axons (eYFP mice: 578.6 ± 36.02 μm; trkB.DN mice: 591.5 ± 40.22μm; *p* = 0.9334). (**g-i**) Scholl profile, the total length, and total number of intersections for the dendrites. (**g**) Primary dendrites: total length; eYFP mice: 1851 ± 139.6 μm; trkB.DN mice: 2003 ± 200.5 μm; *p* = 0.5331, total number of intersections; eYFP mice: 153.9 ± 11.8 intersections; trkB.DN mice: 157.6 ± 17.5 intersections; *p* = 0.8611. (**h**) Secondary dendrites: total length; eYFP mice: 1663 ± 95.9 μm; trkB.DN mice: 2252 ± 206.5 μm; *p* = 0.0138, total number of intersections: eYFP mice: 130.1 ± 7.5 intersections; trkB.DN mice: 166.7 ± 17.5 intersections; *p* = 0.0598. (**i**) Tertiary dendrites: total length; eYFP mice: 407.6 ± 75.0 μm; trkB.DN mice: 811.2 ± 141.8 μm; *p* = 0.0166, total number of intersections; eYFP mice: 31.6 ± 5.6 intersections; trkB.DN mice: 58.1 ± 8.6 intersections; *p* = 0.0147. (**j-m**) Scholl profile, the total length, and total number of intersections for the axons. (**j**) Primary axons: total length; eYFP mice: 634.4 ± 193.8; trkB.DN mice: 1196 ± 270.4; *p* = 0.1546), total number of intersections; eYFP mice: 190.4 ± 16.8; trkB.DN mice: 275.2 ± 42.5; *p* = 0.0685. (**k**) Secondary axons: total length; eYFP mice: 2244 ± 198.8; trkB.DN mice: 3182 ± 497.3; *p* = 0.0840), total number of intersections; eYFP mice: 163.6 ± 15.5; trkB.DN mice: 219.7 ± 29.9; *p* = 0.1010. (**I**) Sholl profile of tertiary and > tertiary axons. Blue shading: significant difference between trkB.DN and eYFP mice. (**m**) Left: tertiary axons: total length; eYFP mice: 5421 ± 554.3; trkB.DN mice: 7042 ± 479.3; *p* = 0.0375, total number of intersections; eYFP mice: 371.0 ± 38.7; trkB.DN mice: 450.4 ± 30.6; *p* = 0.1241. Right: > tertiary axons: total length; eYFP mice: 16030 ± 1450; trkB.DN mice: 23184 ± 1745; *p* = 0.0040, total number of intersections; eYFP mice: 1041 ± 91.4; trkB.DN mice: 1436 ± 114.2; *p* = 0.0118. (**n**) Membrane capacitance of mPFC PV interneurons in trkB.DN mice (82.6 pF) and eYFP mice (63.1 pF); *p* = 0.0009) (**o**) Input resistance of mPFC PV interneurons in trkB.DN mice (103.68 MΩ), and eYFP mice (112.81 MΩ); *p* = 0.4). Data shown as mean ± SEM. For boxplots (n, o), data shown as median, box: 25th and 75th percentile and whiskers: data points that are not outliers. For Sholl analysis plot (g-l), significance was tested with Multiple t tests with the Holm-Sidak method. For bar plots, Unpaired t test, Two-tailed was used to assess significance if data passed the D’Agostino & Pearson normality test, if not Wilcoxon rank-sum test was used. * *p* < 0.05, ** *p* < 0.01, *** *p* < 0.001.

**Figure S3.**
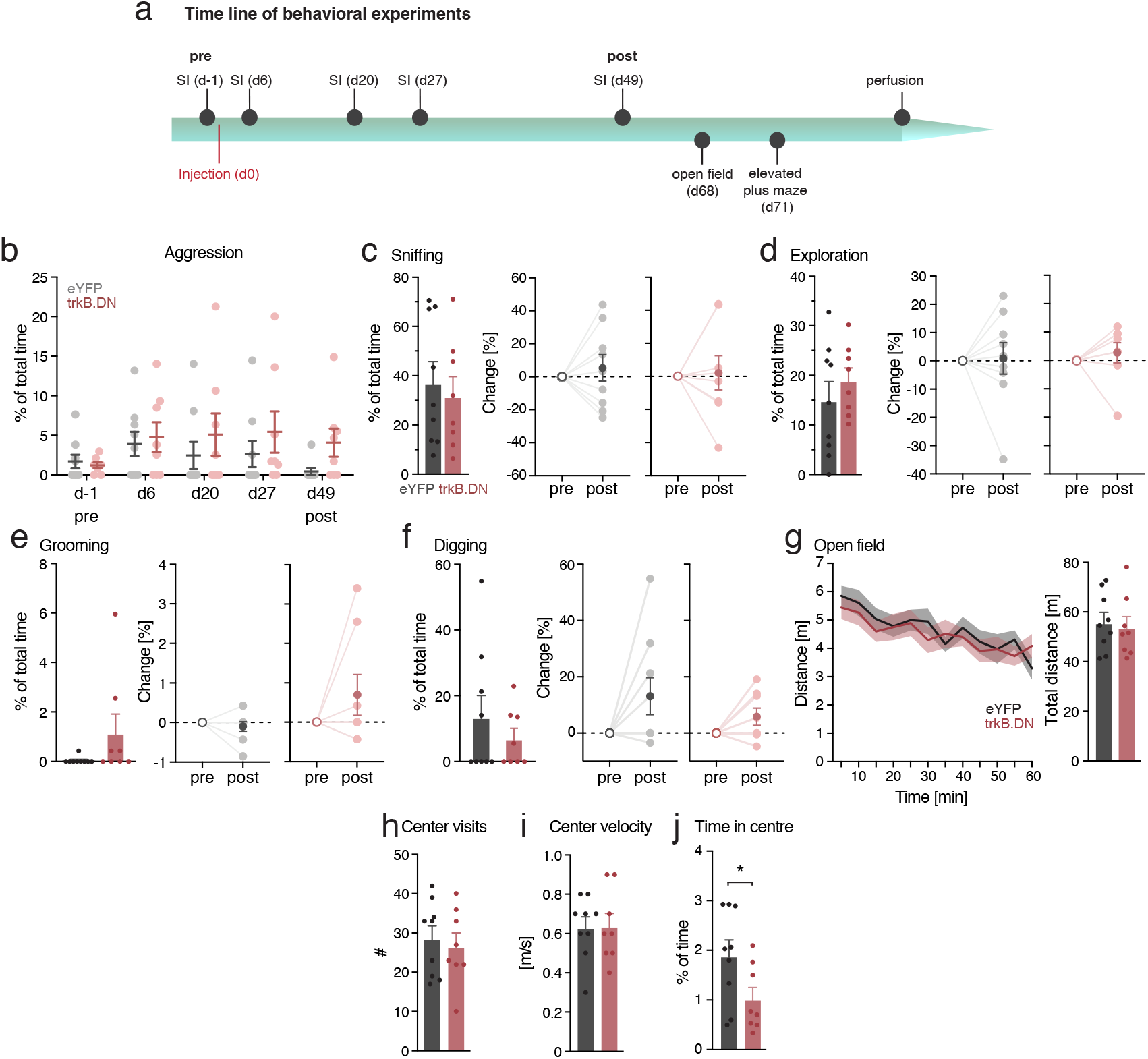
Behavior. (**a**) Timeline of behavioral experiments. (**b**) Percentage of time in aggression in the resident intruder procedure one day before viral injection (d-1; pre), and 6, 20, 27 and 49 (d49; post) days after injection for trkB.DN (n = 8 males), and eYFP mice (n = 9 males). (**c-f**) Non-aggressive behaviors scored in the resident intruder procedure. (No differences were found between trkB.DN and eYFP mice except for measures of aggression). Left: d49 (post), right: change between timepoints pre (d-1) and post (d49). (**c**) Left: percent time in sniffing d49 (post): eYFP mice: 37.1 ± 8.6%; trkB.DN mice: 31.9 ± 7.8%, *p* = 0.6612). Right: Change (percent of time) in sniffing (pre to post): eYFP mice: 5.0 ± 8.2 %, *p* = 0.3672; trkB.DN mice: 2.2 ± 10.5 %; *p* = 0.4727. (**d**) Left: percent time in exploration d49 (post): eYFP mice: 15 ± 3.8%; trkB.DN mice: 19 ± 2.5%, *p* = 0.3960. Right: Change (percent of time) in exploration (pre to post): eYFP mice: 0.9 ± 5.6 %, *p* = 0.3672; trkB.DN mice: 2.9 ± 3.6 %, *p* = 0.1172. (**e**) Left: percent time in grooming d49 (post): eYFP mice: 0.05 ± 0.05 %; trkB.DN mice: 1.17 ± 0.749%, *p* = 0.0905). Right: Change (percent of time) in grooming (pre to post): eYFP mice: −0.09 ± 0.12 %, *p* = 0.375), trkB.DN mice: 0.69 ± 0.51, *p* = 0.25. (**f**) Left: percent time in digging d49 (post): eYFP mice: 13.6 ± 6.5%; trkB.DN mice: 7 ± 3.1%, *p* =0.5812. Right: Change (percent of time) in digging (pre to post): eYFP mice: 13.10 ± 6.59%, *p* = 0.0625; trkB.DN mice: 5.851 ± 3.038 %, *p* = 0.0781. (**g-j**) Open field test. Same mice as in (a-f), performed d68. (**g**) No difference was found in locomotion between trkB.DN mice and eYFP mice. Locomotion during 60 min: eYFP mice: 55.94 ± 3.87; trkB.DN mice: 53.83 ± 4.39 meters, *p* = 0.7229. Bin size: 5 min. (**h**) No difference was found in the number of center visits the first 3 minutes of the open field test between trkB.DN and eYFP mice: eYFP mice: 28.7 ± 3.2; trkB.DN mice: 26.6 ± 3.4 center visits; *p* = 0.666. (**i**) No difference was found in the velocity in the center of the open field box for the first 3 minutes of the test: eYFP mice: 0.63 ± 0.05; trkB.DN mice: 0.64 ± 0.07 m/s, *p* = 0.9607. (**j**) TrkB.DN mice spent significantly less time in the center of the open field box during the first 3 minutes than eYFP mice: eYFP: 1.90 ± 0.31%; trkB.DN: 1.03 ± 0.23% of time; *p* = 0.0458. Data shown as mean ± SEM. For bar plots, Unpaired t test - two-tailed was used to assess significance if data passed the D’Agostino & Pearson normality test, if not Wilcoxon rank-sum test was used. For (g) 2way ANOVA, Sidak’s multiple comparisons test was used. For paired data, a Wilcoxon matched pairs test was used. * *p* < 0.05

**Figure S4.**
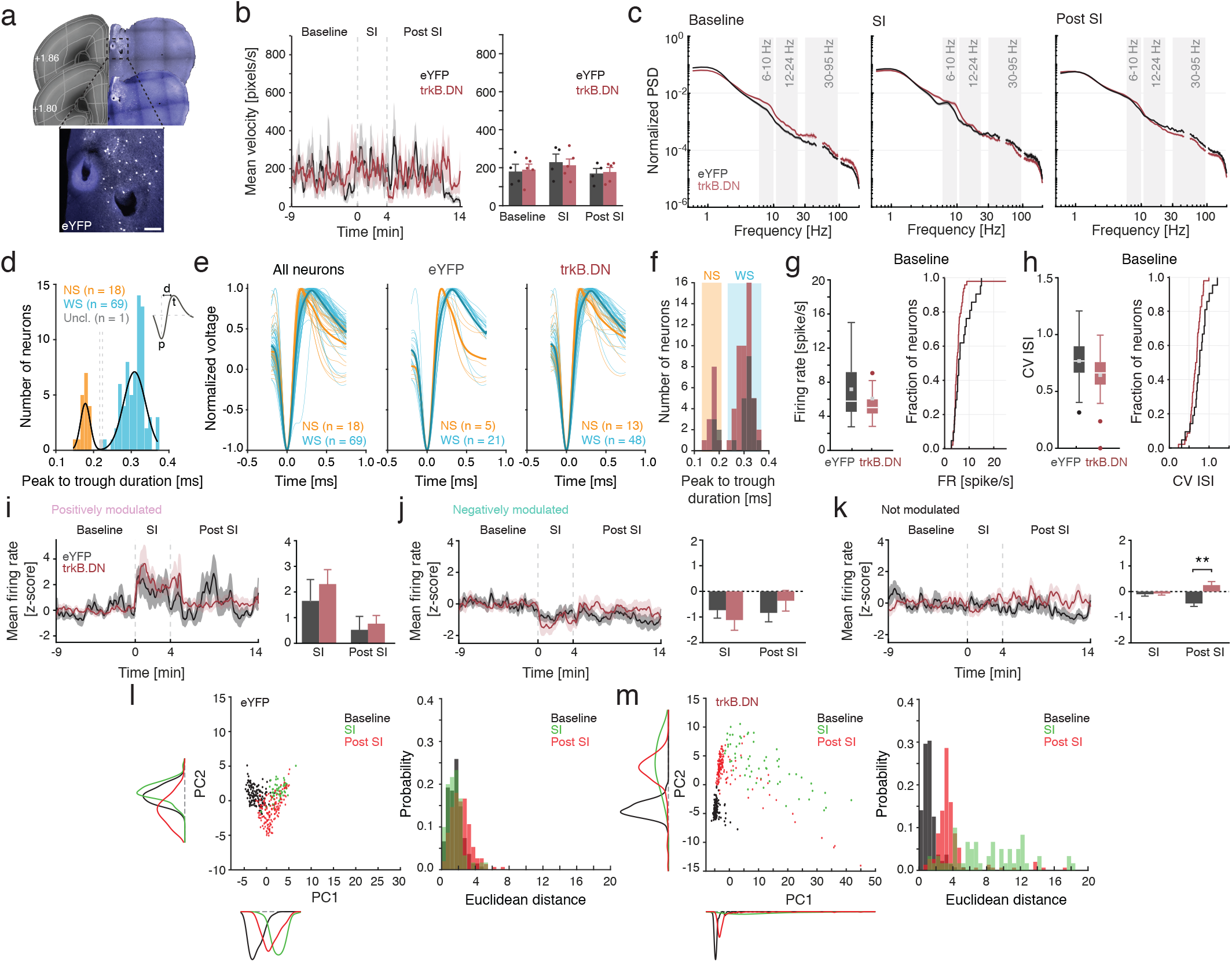
Social interaction recordings. (**a-m**) Chronic extracellular single-unit and LFP recordings during social interaction. eYFP mice (gray; n = 4 males), trkB.DN mice (red; n = 5 males). (**a**) Example brain sections from an eYFP mouse with electrolytic lesions used for reconstruction of the tetrodes positions (related to Fig. 4a). (**b**) There is no difference in the mean velocity of locomotion for implanted eYFP mice and trkB.DN mice during baseline, SI, or Post SI: baseline: eYFP mice: 179.8 ± 38.5; trkB.DN mice: 189.8 ± 29.1 pixels/s, *p* = 0.839, SI: eYFP mice: 229.3 ± 42.5; trkB.DN mice: 212.8 ± 32.9 pixels/s, *p* = 0.764, Post SI: eYFP mice: 169.5 ± 26.5; trkB.DN mice: 177.3 ± 24.1 pixels/s, *p* = 0.834. (**c**) Direct comparison of bootstrapped normalized PSD of the LFP for baseline, SI, and Post SI between eYFP (**gray**) and trkB.DN mice (**red**). (**d**) Classification of recorded single units into narrow-spiking (NS) putative inhibitory interneurons and wide-spiking (WS) putative excitatory pyramidal neurons based on spike waveform features (the peak to trough duration (d). NS (orange): n = 18, mean d = 0.177 ± 0.012 ms; WS (light blue): n = 69, mean d = 0.309 ± 0.028 ms; gray: unclassified units (n =1). Inlet: The spike waveform features used to characterize action potentials: peak (p), trough (t), and peak to trough duration (d). (**e**) Normalized average spike waveforms of the classified NS (orange) and WS (light blue) neurons. Left: All neurons; middle: eYFP mice; right; trkB.DN mice. (**f**) The spike waveforms (peak to trough duration) of NS (orange shading) and WS (light blue shading) neurons do not differ between trkB.DN (red) and eYFP mice (gray): NS: eYFP mice: 0.183 ± 0.009 ms; trkB.DN mice: 0.175 ± 0.013 ms, *p* = 0.057), WS: eYFP mice: 0.319 ± 0.025 ms; trkB.DN mice: 0.305 ± 0.029 ms, *p* = 0.22. (**g-m**) Activity of WS neurons. (**g-h**) There is no difference in the baseline (−9-0 min) firing of WS neurons between trkB.DN (red) and eYFP (gray) mice. (**g**) Left: boxplots of the average firing rate of WS neurons during baseline: eYFP mice: 7.2 ± 0.17; trkB.DN mice: 6.08 ± 0.12 spike/s. Right: the cumulative distribution of the firing rate of WS neurons during baseline; *p* = 0.09. (**h**) Left: boxplots of the coefficient of variation of the inter-spike interval (CV ISI) distribution during baseline: eYFP mice: 0.76 ± 0.01; trkB.DN mice: 0.637 ± 0.004. Right: the cumulative distribution of the CV ISI during baseline, *p* = 0.08. (**i-k**) Subdivision of the WS population into positively, negatively, and not modulated subpopulations based on the firing rate of individual neurons during SI (0-4 min). Related to Fig. 4j. (**i**) Mean firing rate (z-scored) of positively modulated WS neuron: SI: eYFP mice: 1.65 ± 0.83; trkB.DN mice: 2.31 ± 0.56; *p* = 0.72, Post SI: eYFP mice: 0.52 ± 0.52; trkB.DN mice: 0.76 ± 0.32, *p* = 0.12. (**j**) Mean firing rate (z-scored) of negatively modulated WS neuron: SI: eYFP mice: −0.73 ± 0.16; trkB.DN mice: −1.12 ± 0.32, *p* = 0.66), Post SI: eYFP mice: −0.84 ± 0.20; trkB.DN mice: −0.37 ± 0.30, *p* = 0.28. (**k**) Mean firing rate (z-scored) of not modulated WS neuron: SI: eYFP mice: −0.11 ± 0.07; trkB.DN mice: −0.08 ± 0.06, *p* = 1.00); Post SI: eYFP mice: −0.46 ± 0.12; trkB.DN mice: 0.26 ± 0.14; *p* = 0.002 (**l-m**) Left: Scatter plot and kernel density of 2D projection of population dynamics. Each data point corresponds to the first and second PC calculated over a bin of 4 s. Clusters were defined according to the period of experiment (Baseline, black; SI, green; and Post SI, red). Right: Probability histogram of intra-cluster Euclidean Distances. (**I**) Population dynamics representation (left) resulted in overlapping but well-defined clusters for baseline, SI and Post SI in eYFP mice. Euclidean distance (right) differs between clusters after social interaction (One-way ANOVA, F_cluster(2,342)_ = 19.1, *p* <0.001). Euclidean distance was similar between Baseline and SI (*p* = 1), but different between SI and Post SI (*p* < 0.001) and between Baseline and Post SI (*p* < 0.001). (**m**) Population dynamics representation in trkB.DN mice (left) resulted in non-overlapping and not well-defined clusters, with specific increased variability along the PC1 axis. Euclidean distance probability (right) differs between clusters (One-way ANOVA, F_cluster(2,342)_ = 52.7, *p* < 0.001). A Bonferroni post hoc revealed differences in Euclidean distance between Baseline and SI (*p* < 0.001), SI and Post SI (*p* < 0.001) and between Baseline and Post SI (*p* < 0.001). Data shown as mean ± SEM for (b, c and i-k). For boxplots (g-h), data shown as median (white line), mean (squared dot), box: 25th and 75th percentile and whiskers: data points that are not outliers, dots: outliers. For bar plots, Unpaired t test - two-tailed was used to assess significance if data passed the Kolmogorov-Smirnov normality test, if not the Wilcoxon test was used. For cumulative distribution function (g-h), the Kolmolgorov-Smirnov test was used. For (l-m) One-way ANOVA was used along with a Bonferroni post hoc test. ** *p* < 0.01.

**Figure S5.**
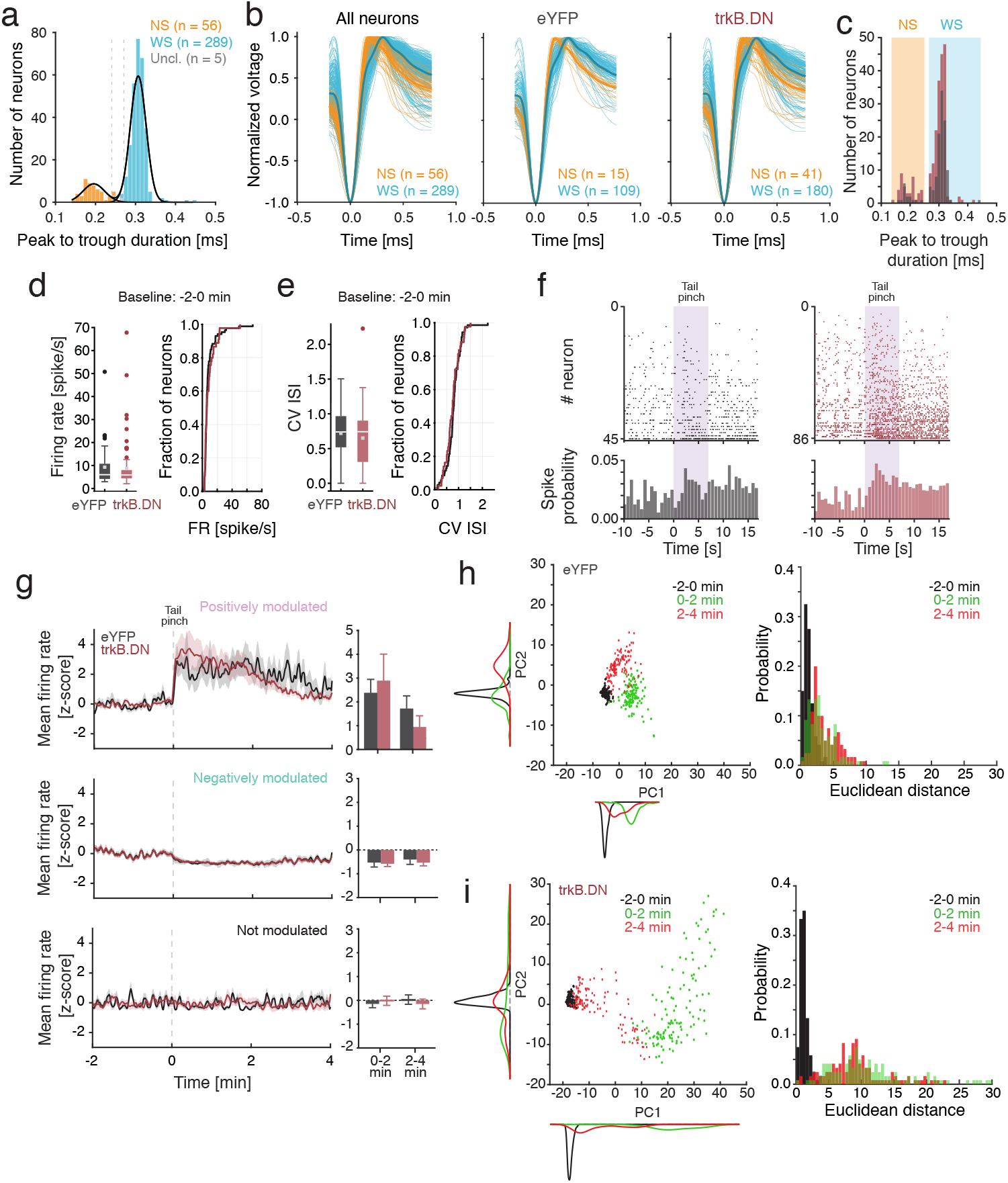
Tail pinch. (**a**) Classification of recorded single units into narrow-spiking (NS) putative inhibitory interneurons and wide-spiking (WS) putative excitatory pyramidal neurons based on spike waveform features (the peak to trough duration (d)). NS (orange): n = 56, mean d = 0.194 ± 0.025 ms; WS (light blue): n = 289, mean d = 0.307 ± 0.019 ms; gray: unclassified units (n =5). (**b**) Normalized average spike waveforms of the classified NS (orange) and WS (light blue) neurons. Left: All neurons; middle: eYFP mice; right; trkB.DN mice. (**c**) The spike waveforms (peak to trough duration) of NS (orange shading) and WS (light blue shading) neurons do not differ between trkB.DN (red) and eYFP mice (gray): NS: eYFP mice: 0.191 ± 0.021 ms; trkB.DN mice: 0.195 ± 0.027 ms; *p* = 0.22, WS: eYFP mice: 0.309 ± 0.019 ms; trkB.DN mice: 0.306 ± 0.019 ms; *p* = 0.79. (**d-h**) Activity of WS neurons. (**d-e**) There is no difference in the baseline (−2-0 min) firing of WS neurons between trkB.DN (red) and eYFP (gray) mice. (**d**) Left: boxplots of the distribution of the average firing rate of WS neurons during baseline: eYFP mice: 9.28 ± 0.19; trkB.DN mice: 8.67 ± 0.11 spike/s. Right: the cumulative distribution of the firing rate of WS neurons during baseline; *p* = 0.58. (**e**) Left: boxplots of the coefficient of variation of the inter-spike interval (CV ISI) distribution during baseline: eYFP mice: 0.72 ± 0.01; trkB.DN mice: 0.65 ± 0.01. Right: the cumulative distribution of the CV ISI during baseline, *p* = 0.43. (**f**) Top: raster plot of the spiking of the WS neurons in eYFP mice (gray) and trkB.DN mice (red) around the time of tail pinch (purple bar; 7 s). Bottom: the spike probability. (**g**) Subdivision of the WS population into positively (top), negatively (middle), and not modulated (bottom) subpopulations based on the firing rate of the individual neurons during tail pinch (0-2 min). No differences in the mean firing rate of the WS populations were found for any of the three subpopulations between trkB.DN and eYFP mice. The mean firing rates 0-2 min, and 2-4 min, respectively, were compared. Positively modulated: 0-2 min; eYFP: 2.38 ± 0.38; trkB.DN: 2.89 ±1 .02; *p* = 0.37), 2-4 min; eYFP: 1.72 ± 0.36; trkB.DN: 0.95 ± 0.30; *p* = 0.22. Negatively modulated: eYFP: 0-2 min; −0.13 ± 0.10; trkB.DN: −0.58 ± 0.05; *p* = 0.15, 2-4 min; eYFP: −0.40 ± 0.14; trkB.DN: −0.53 ± 0.06; *p* = 0.06. Not modulated: 0–2 min; eYFP: −0.15 ± 0.02; trkB.DN: −0.02 ± 0.03; *p* = 0.08, 2–4 min: eYFP: 0.03 ± 0.10; trkB.DN: −0.15 ± 0.10; *p* = 0.79. Dashed line: start of tail pinch. Related to Fig. 5f. **h-i**) Left: Scatter plot and kernel density of 2D projection of population dynamics. Each data point corresponds to the first and second PC calculated over a bin of 4 s. Clusters were defined according to the period of experiment around tail pinch (−2 - 0 min, black; 0 - 2 min, green; and 2 - 4 min, red). Right: Probability histogram of intra-cluster Euclidean Distances. (**h**) Population dynamics representation (left) resulted in well-defined clusters for baseline, SI and Post SI in eYFP mice. Euclidean distance (right) differs between clusters (One-way ANOVA, F_cluster(2,357)_ = 69.4, *p* <0.001). A Bonferroni post hoc revealed differences in Euclidean distance between Baseline and 0-2 min after tail pinch (*p* < 0.001), between 0-2 min and 2-4 min after tail pinch (*p* = 0.04), and between Baseline and 2-4 min (*p* < 0.001). (**i**) Population dynamics representation in trkB.DN mice (left) resulted in dispersed and not well-defined clusters, with specific increased variability along the PC1 axis. Euclidean distance (right) differs between clusters (One-way ANOVA, F_cluster(2,357)_ = 167.8, *p* <0.001). Differences in Euclidean distance were detected between Baseline and 0-2 min after tail pinch (*p* < 0.001), between 0-2 min and 2-4 min after tail pinch (*p* = 0.004), and between Baseline and 2-4 min (*p* < 0.001). For boxplots (d-e), data shown as median (white line), mean (squared dot), box: 25th and 75th percentile and whiskers: data points that are not outliers, dots: outliers. For (g) data shown as mean ± SEM. For (g), the Wilcoxon rank-sum test was used to assess significance since data did not pass the Kolmogorov-Smirnov normality test. For (h-i) One-way ANOVA was used along with a Bonferroni post hoc test.

## References

Agetsuma, Masakazu, Jordan P Hamm, Kentaro Tao, Shigeyoshi Fujisawa, and Rafael Yuste. 2018. “Parvalbumin-Positive Interneurons Regulate Neuronal Ensembles in Visual Cortex.” Cerebral Cortex 28 (5): 1831–45. https://doi.org/10.1093/cercor/bhx169.

Ardid, Salva, Martin Vinck, D. Kaping, Susanna Marquez, Stefan Everling, and Thilo Womelsdorf. 2015. “Mapping of Functionally Characterized Cell Classes onto Canonical Circuit Operations in Primate Prefrontal Cortex.” Journal of Neuroscience 35 (7): 2975–91. https://doi.org/10.1523/JNEUROSCI.2700-14.2015.

Armanini, M. P., S. B. McMahon, J. Sutherland, D. L. Shelton, and H. S. Phillips. 1995. “Truncated and Catalytic Isoforms of TrkB Are Co‐expressed in Neurons of Rat and Mouse CNS.” European Journal of Neuroscience 7 (6): 1403–9. https://doi.org/10.1111/j.1460-9568.1995.tb01132.x.

Bakker, Rembrandt, Paul Tiesinga, and Rolf Kötter. 2015. “The Scalable Brain Atlas: Instant Web-Based Access to Public Brain Atlases and Related Content.” https://doi.org/10.1007/s12021-014-9258-x.

Barfield, Elizabeth T., and Shannon L. Gourley. 2018. “Prefrontal Cortical TrkB, Glucocorticoids, and Their Interactions in Stress and Developmental Contexts.” Neuroscience & Biobehavioral Reviews 95 (December): 535–58. https://doi.org/10.1016/j.neubiorev.2018.10.015.

Bartos, Marlene, Imre Vida, and Peter Jonas. 2007. “Synaptic Mechanisms of Synchronized Gamma Oscillations in Inhibitory Interneuron Networks.” Nature Reviews Neuroscience 8 (1): 45–56. https://doi.org/10.1038/nrn2044.

Baxter, Gregory T., Monte J. Radeke, Richard C. Kuo, Victoria Makrides, Beth Hinkle, Richard Hoang, Angelica Medina-Selby, Doris Coit, Pablo Valenzuela, and Stuart C. Feinstein. 1997. “Signal Transduction Mediated by the Truncated TrkB Receptor Isoforms, TrkB.T1 and TrkB.T2.” Journal of Neuroscience 17 (8): 2683–90. https://doi.org/10.1523/jneurosci.17-08-02683.1997.

Berg, Stuart, Dominik Kutra, Thorben Kroeger, Christoph N. Straehle, Bernhard X. Kausler, Carsten Haubold, Martin Schiegg, et al. 2019. “Ilastik: Interactive Machine Learning for (Bio)Image Analysis.” Nature Methods 16 (12): 1226–32. https://doi.org/10.1038/s41592-019-0582-9.

Bracken, Bethany K., and Gina G. Turrigiano. 2009. “Experience-Dependent Regulation of TrkB Isoforms in Rodent Visual Cortex.” Developmental Neurobiology 69 (5): 267–78. https://doi.org/10.1002/dneu.20701.

Butler, Andrew, Paul Hoffman, Peter Smibert, Efthymia Papalexi, and Rahul Satija. 2018. “Integrating Single-Cell Transcriptomic Data across Different Conditions, Technologies, and Species.” Nature Biotechnology 36 (5): 411–20. https://doi.org/10.1038/nbt.4096.

Buzsáki, György, and Xiao-Jing Wang. 2012. “Mechanisms of Gamma Oscillations.” Annual Review of Neuroscience 35 (1): 203–25. https://doi.org/10.1146/annurev-neuro-062111-150444.

Cardin, Jessica A., Marie Carlén, Konstantinos Meletis, Ulf Knoblich, Feng Zhang, Karl Deisseroth, Li-Huei Tsai, and Christopher I Moore. 2009. “Driving Fast-Spiking Cells Induces Gamma Rhythm and Controls Sensory Responses.” Nature 459 (7247): 663–67. https://doi.org/10.1038/nature08002.

Carim-Todd, Laura, Kevin G Bath, Gianluca Fulgenzi, Sudhirkumar Yanpallewar, Deqiang Jing, Colleen a Barrick, Jodi Becker, et al. 2009. “Endogenous Truncated TrkB.T1 Receptor Regulates Neuronal Complexity and TrkB Kinase Receptor Function In Vivo.” Journal of Neuroscience 29 (3): 678–85. https://doi.org/10.1523/JNEUROSCI.5060-08.2009.

Carlén, M, K Meletis, J H Siegle, J a Cardin, K Futai, D Vierling-Claassen, C Rühlmann, et al. 2012. “A Critical Role for NMDA Receptors in Parvalbumin Interneurons for Gamma Rhythm Induction and Behavior.” Molecular Psychiatry 17 (5): 537–48. https://doi.org/10.1038/mp.2011.31.

Cellerino, Alessandro, Lamberto Maffei, and Luciano Domenici. 1996. “The Distribution of Brain-Derived Neurotrophic Factor and Its Receptor TrkB in Parvlbumin-Containing Neurons of the Rat Visual Cortex.” European Journal of Neuroscience 8 (6): 1190–97. https://doi.org/10.1111/j.1460-9568.1996.tb01287.x.

Cho, Kathleen K.A., Renee Hoch, Anthony T. Lee, Tosha Patel, John L.R. Rubenstein, and Vikaas S. Sohal. 2015. “Gamma Rhythms Link Prefrontal Interneuron Dysfunction with Cognitive Inflexibility in Dlx5/6+/− Mice.” Neuron 85 (6): 1332–43. https://doi.org/10.1016/j.neuron.2015.02.019.

Dienel, Samuel J., and David A. Lewis. 2018. “Alterations in Cortical Interneurons and Cognitive Function in Schizophrenia.” Neurobiology of Disease, no. January: 0–1. https://doi.org/10.1016/j.nbd.2018.06.020.

Eide, F F, E R Vining, B L Eide, K Zang, X Y Wang, and L F Reichardt. 1996. “Naturally Occurring Truncated TrkB Receptors Have Dominant Inhibitory Effects on Brain-Derived Neurotrophic Factor Signaling.” The Journal of Neuroscience : The Official Journal of the Society for Neuroscience 16 (10): 3123–29. https://doi.org/10.1055/s-0029-1237430.Imprinting.

Fenner, Barbara M. 2012. “Truncated TrkB: Beyond a Dominant Negative Receptor.” Cytokine & Growth Factor Reviews 23 (1–2): 15–24. https://doi.org/10.1016/j.cytogfr.2012.01.002.

Ferguson, Brielle R., and Wen-Jun Gao. 2018. “PV Interneurons: Critical Regulators of E/I Balance for Prefrontal Cortex-Dependent Behavior and Psychiatric Disorders.” Frontiers in Neural Circuits 12 (May): 37. https://doi.org/10.3389/fncir.2018.00037.

Ferreira, Tiago A., Arne V. Blackman, Julia Oyrer, Sriram Jayabal, Andrew J. Chung, Alanna J. Watt, P. Jesper Sjöström, and Donald J. Van Meyel. 2014. “Neuronal Morphometry Directly from Bitmap Images.” Nature Methods. Nature Publishing Group. https://doi.org/10.1038/nmeth.3125.

Gorba, T, and P Wahle. 1999. “Expression of TrkB and TrkC but Not BDNF MRNA in Neurochemically Identified Interneurons in Rat Visual Cortex in Vivo and in Organotypic Cultures.” The European Journal of Neuroscience 11 (4): 1179–90.

Grech, Adrienne Mary, Xin Du, Simon S. Murray, Junhua Xiao, and Rachel Anne Hill. 2019. “Sex-Specific Spatial Memory Deficits in Mice with a Conditional TrkB Deletion on Parvalbumin Interneurons.” Behavioural Brain Research 372 (October): 111984. https://doi.org/10.1016/j.bbr.2019.111984.

Hamm, Jordan P., Darcy S. Peterka, Joseph A. Gogos, and Rafael Yuste. 2017. “Altered Cortical Ensembles in Mouse Models of Schizophrenia.” Neuron 94 (1): 153–167.e8. https://doi.org/10.1016/j.neuron.2017.03.019.

Hashimoto, Takanori, Sarah E Bergen, Quyen L Nguyen, Baoji Xu, Lisa M Monteggia, Joseph N Pierri, Zhuoxin Sun, Allan R Sampson, and David a Lewis. 2005. “Relationship of Brain-Derived Neurotrophic Factor and Its Receptor TrkB to Altered Inhibitory Prefrontal Circuitry in Schizophrenia.” The Journal of Neuroscience : The Official Journal of the Society for Neuroscience 25 (2): 372–83. https://doi.org/10.1523/JNEUROSCI.4035-04.2005.

Holtmaat, Anthony, and Pico Caroni. 2016. “Functional and Structural Underpinnings of Neuronal Assembly Formation in Learning.” Nature Neuroscience 19 (12): 1553–62. https://doi.org/10.1038/nn.4418.

Hu, Hua, Jian Gan, and Peter Jonas. 2014. “Fast-Spiking, Parvalbumin+ GABAergic Interneurons: From Cellular Design to Microcircuit Function.” Science 345 (6196): 1255263–1255263. https://doi.org/10.1126/science.1255263.

Huang, Z J, A Kirkwood, T Pizzorusso, V Porciatti, B Morales, M F Bear, L Maffei, and S Tonegawa. 1999. “BDNF Regulates the Maturation of Inhibition and the Critical Period of Plasticity in Mouse Visual Cortex.” Cell 98 (6): 739–55. https://doi.org/10.1016/s0092-8674(00)81509-3.

Ilchibaeva, Tatiana V., Anton S. Tsybko, Rimma V. Kozhemyakina, Elena M. Kondaurova, Nina K. Popova, and Vladimir S. Naumenko. 2018. “Genetically Defined Fear-Induced Aggression: Focus on BDNF and Its Receptors.” Behavioural Brain Research 343 (May): 102–10. https://doi.org/10.1016/j.bbr.2018.01.034.

Jadi, Monika P., M. Margarita Behrens, and Terrence J. Sejnowski. 2016. Abnormal Gamma Oscillations in N-Methyl-D-Aspartate Receptor Hypofunction Models of Schizophrenia. Biological Psychiatry. Vol. 79. https://doi.org/10.1016/j.biopsych.2015.07.005.

Jiao, Yuanyuan, Z. Zhang, Chunzhao Zhang, Xinjun Wang, Kazuko Sakata, Bai Lu, and Q.-Q. Sun. 2011. “A Key Mechanism Underlying Sensory Experience-Dependent Maturation of Neocortical GABAergic Circuits in Vivo.” Proceedings of the National Academy of Sciences 108 (29): 12131–36. https://doi.org/10.1073/pnas.1105296108.

Korotkova, Tatiana, Elke C. Fuchs, Alexey Ponomarenko, Jakob von Engelhardt, and Hannah Monyer. 2010. “NMDA Receptor Ablation on Parvalbumin-Positive Interneurons Impairs Hippocampal Synchrony, Spatial Representations, and Working Memory.” Neuron 68 (3): 557–69. https://doi.org/10.1016/j.neuron.2010.09.017.

Kowiański, Przemysław, Grażyna Lietzau, Ewelina Czuba, Monika Waśkow, Aleksandra Steliga, and Janusz Moryś. 2018. “BDNF: A Key Factor with Multipotent Impact on Brain Signaling and Synaptic Plasticity.” Cellular and Molecular Neurobiology. Springer New York LLC. https://doi.org/10.1007/s10571-017-0510-4.

Levy, Dana Rubi, Tal Tamir, Maya Kaufman, Ana Parabucki, Aharon Weissbrod, Elad Schneidman, and Ofer Yizhar. 2019. “Dynamics of Social Representation in the Mouse Prefrontal Cortex.” Nature Neuroscience 22 (12): 2013–22. https://doi.org/10.1038/s41593-019-0531-z.

Longair, Mark H, Dean A Baker, and J Douglas Armstrong. 2011. “Simple Neurite Tracer: Open Source Software for Reconstruction, Visualization and Analysis of Neuronal Processes.” Bioinformatics 27 (17): 2453–54. https://doi.org/10.1093/bioinformatics/btr390.

Luberg, Kristi, Jenny Wong, Cynthia Shannon Weickert, and Tõnis Timmusk. 2010. “Human TrkB Gene: Novel Alternative Transcripts, Protein Isoforms and Expression Pattern in the Prefrontal Cerebral Cortex during Postnatal Development.” Journal of Neurochemistry 113 (4): 952–64. https://doi.org/10.1111/j.1471-4159.2010.06662.x.

Lucas, Elizabeth K, Anita Jegarl, and Roger L Clem. 2014. “Mice Lacking TrkB in Parvalbumin-Positive Cells Exhibit Sexually Dimorphic Behavioral Phenotypes.” Behavioural Brain Research, August, 1–7. https://doi.org/10.1016/j.bbr.2014.08.011.

Mantz, J, C Milla, J Glowinski, and A M Thierry. 1988. “Differential Effects of Ascending Neurons Containing Dopamine and Noradrenaline in the Control of Spontaneous Activity and of Evoked Responses in the Rat Prefrontal Cortex.” Neuroscience 27 (2): 517–26. https://doi.org/10.1016/0306-4522(88)90285-0.

Märtin, Antje, Daniela Calvigioni, Ourania Tzortzi, Janos Fuzik, Emil Wärnberg, and Konstantinos Meletis. 2019. “A Spatiomolecular Map of the Striatum.” Cell Reports 29 (13): 4320–4333.e5. https://doi.org/10.1016/j.celrep.2019.11.096.

Massi, Lema, Michael Lagler, Katja Hartwich, Zsolt Borhegyi, Peter Somogyi, and Thomas Klausberger. 2012. “Temporal Dynamics of Parvalbumin-Expressing Axo-Axonic and Basket Cells in the Rat Medial Prefrontal Cortex in Vivo.” Journal of Neuroscience 32 (46): 16496–502. https://doi.org/10.1523/JNEUROSCI.3475-12.2012.

Mathalon, Daniel H., and Vikaas S. Sohal. 2015. “Neural Oscillations and Synchrony in Brain Dysfunction and Neuropsychiatric Disorders It’s about Time.” JAMA Psychiatry 72 (8): 840–44. https://doi.org/10.1001/jamapsychiatry.2015.0483.

Mathis, Alexander, Pranav Mamidanna, Kevin M. Cury, Taiga Abe, Venkatesh N. Murthy, Mackenzie Weygandt Mathis, and Matthias Bethge. 2018. “DeepLabCut: Markerless Pose Estimation of User-Defined Body Parts with Deep Learning.” Nature Neuroscience 21 (9): 1281–89. https://doi.org/10.1038/s41593-018-0209-y.

Mikics, Éva, Ramon Guirado, Juzoh Umemori, Máté Tóth, László Biró, Christina Miskolczi, Diána Balázsfi, et al. 2018. “Social Learning Requires Plasticity Enhanced by Fluoxetine through Prefrontal Bdnf-TrkB Signaling to Limit Aggression Induced by Post-Weaning Social Isolation.” Neuropsychopharmacology 43 (2): 235–45. https://doi.org/10.1038/npp.2017.142.

Moore, Christopher I., Marie Carlen, Ulf Knoblich, and Jessica a. Cardin. 2010. “Neocortical Interneurons: From Diversity, Strength.” Cell 142 (2): 189–93. https://doi.org/10.1016/j.cell.2010.07.005.

Nagappan, Guhan, and Bai Lu. 2005. “Activity-Dependent Modulation of the BDNF Receptor TrkB: Mechanisms and Implications.” Trends in Neurosciences 28 (9): 464–71. https://doi.org/10.1016/j.tins.2005.07.003.

Notaras, Michael, and Maarten van den Buuse. 2020. “Neurobiology of BDNF in Fear Memory, Sensitivity to Stress, and Stress-Related Disorders.” Molecular Psychiatry, January. https://doi.org/10.1038/s41380-019-0639-2.

O’Neill, Kate M., Barbara F. Akum, Survandita T. Dhawan, Munjin Kwon, Christopher G. Langhammer, and Bonnie L. Firestein. 2015. “Assessing Effects on Dendritic Arborization Using Novel Sholl Analyses.” Frontiers in Cellular Neuroscience 9 (JULY). https://doi.org/10.3389/fncel.2015.00285.

Ohira, K., and M. Hayashi. 2009. “A New Aspect of the TrkB Signaling Pathway in Neural Plasticity.” Current Neuropharmacology 7 (4): 276–85. https://doi.org/10.2174/157015909790031210.

Öner, Metin, and Ipek Deveci Kocakoç. 2017. “JMASM 49: A Compilation of Some Popular Goodness of Fit Tests for Normal Distribution: Their Algorithms and MATLAB Codes (MATLAB).” Journal of Modern Applied Statistical Methods 16 (2): 547–75. https://doi.org/10.22237/jmasm/1509496200.

Park, Hyungju, and Mu Ming Poo. 2013. “Neurotrophin Regulation of Neural Circuit Development and Function.” Nature Reviews Neuroscience. Nature Publishing Group. https://doi.org/10.1038/nrn3379.

Pessoa, Luiz. 2019. “Neural Dynamics of Emotion and Cognition: From Trajectories to Underlying Neural Geometry.” Neural Networks 120 (December): 158–66. https://doi.org/10.1016/j.neunet.2019.08.007.

Picelli, Simone, Åsa K. Björklund, Omid R. Faridani, Sven Sagasser, Gösta Winberg, and Rickard Sandberg. 2013. “Smart-Seq2 for Sensitive Full-Length Transcriptome Profiling in Single Cells.” Nature Methods 10 (11): 1096–1100. https://doi.org/10.1038/nmeth.2639.

Rodgers, R. J., and A. Dalvi. 1997. “Anxiety, Defence and the Elevated plus-Maze.” In Neuroscience and Biobehavioral Reviews, 21:801–10. Neurosci Biobehav Rev. https://doi.org/10.1016/S0149-7634(96)00058-9.

Schindelin, Johannes, Ignacio Arganda-Carreras, Erwin Frise, Verena Kaynig, Mark Longair, Tobias Pietzsch, Stephan Preibisch, et al. 2012. “Fiji: An Open-Source Platform for Biological-Image Analysis.” Nature Methods. https://doi.org/10.1038/nmeth.2019.

Schmitzer-Torbert, N., J. Jackson, D. Henze, K. Harris, and A. D. Redish. 2005. “Quantitative Measures of Cluster Quality for Use in Extracellular Recordings.” Neuroscience 131 (1): 1–11. https://doi.org/10.1016/j.neuroscience.2004.09.066.

Shamash, Philip, Matteo Carandini, Kenneth Harris, and Nick Steinmetz. 2018. “A Tool for Analyzing Electrode Tracks from Slice Histology.” BioRxiv, October, 447995. https://doi.org/10.1101/447995.

Sohal, Vikaas S., and John L.R. Rubenstein. 2019. “Excitation-Inhibition Balance as a Framework for Investigating Mechanisms in Neuropsychiatric Disorders.” Molecular Psychiatry 24 (9): 1248–57. https://doi.org/10.1038/s41380-019-0426-0.

Sohal, Vikaas S, Feng Zhang, Ofer Yizhar, and Karl Deisseroth. 2009. “Parvalbumin Neurons and Gamma Rhythms Enhance Cortical Circuit Performance.” Nature 459 (7247): 698–702. https://doi.org/10.1038/nature07991.

Stoilov, Peter, Eero Castren, and Stefan Stamm. 2002. “Analysis of the Human TrkB Gene Genomic Organization Reveals Novel TrkB Isoforms, Unusual Gene Length, and Splicing Mechanism.” Biochemical and Biophysical Research Communications 290 (3): 1054–65. https://doi.org/10.1006/bbrc.2001.6301.

Susaki, Etsuo A., Kazuki Tainaka, Dimitri Perrin, Fumiaki Kishino, Takehiro Tawara, Tomonobu M. Watanabe, Chihiro Yokoyama, et al. 2014. “Whole-Brain Imaging with Single-Cell Resolution Using Chemical Cocktails and Computational Analysis.” Cell 157 (3): 726–39. https://doi.org/10.1016/j.cell.2014.03.042.

Tan, Shawn, Yixin Xiao, Henry H Yin, Albert I Chen, Tuck Wah, H Shawn Je, David E. Pozen, Tuck Wah Soong, and H Shawn Je. 2018. “Postnatal TrkB Ablation in Corticolimbic Interneurons Induces Social Dominance in Male Mice.” Proceedings of the National Academy of Sciences 115 (42): E9909–15. https://doi.org/10.1073/pnas.1812083115.

Tremblay, Robin, Soohyun Lee, and Bernardo Rudy. 2016. “GABAergic Interneurons in the Neocortex: From Cellular Properties to Circuits.” Neuron 91 (2): 260–92. https://doi.org/10.1016/j.neuron.2016.06.033.

Voigts, Jakob, Josh Siegle, Dominique L. Pritchett, and Christopher I. Moore. 2013. “The FlexDrive: An Ultra-Light Implant for Optical Control and Highly Parallel Chronic Recording of Neuronal Ensembles in Freely Moving Mice.” Frontiers in Systems Neuroscience 7 (MARCH 2013). https://doi.org/10.3389/fnsys.2013.00008.

Winkel, Frederike, Mathias B Voigt, Giuliano Didio, Salomé Matéo, Elias Jetsonen, Maria Llach Pou, Anna Steinzeig, et al. 2020. “Optical TrkB Activation in Parvalbumin Interneurons Regulates Intrinsic States to Orchestrate Cortical Plasticity.” BioRxiv, January, 2020.04.27.063503. https://doi.org/10.1101/2020.04.27.063503.

Winslow, James T. 2003. “Mouse Social Recognition and Preference.” Current Protocols in Neuroscience 22 (1): 8.16.1–8.16.16. https://doi.org/10.1002/0471142301.ns0816s22.

Wong, Jenny, and Brett Garner. 2012. “Evidence That Truncated TrkB Isoform, TrkB-Shc Can Regulate Phosphorylated TrkB Protein Levels.” Biochemical and Biophysical Research Communications 420 (2): 331–35. https://doi.org/10.1016/j.bbrc.2012.02.159.

Wong, Jenny, Debora A. Rothmond, Maree J. Webster, and Cynthia Shannon Weickert. 2013. “Increases in Two Truncated TrkB Isoforms in the Prefrontal Cortex of People with Schizophrenia.” Schizophrenia Bulletin 39 (1): 130–40. https://doi.org/10.1093/schbul/sbr070.

Woo, Newton H., and Bai Lu. 2006. “Regulation of Cortical Interneurons by Neurotrophins: From Development to Cognitive Disorders.” The Neuroscientist 12 (1): 43–56. https://doi.org/10.1177/1073858405284360.

Xenos, Dionysios, Marija Kamceva, Simone Tomasi, Jessica A. Cardin, Michael L. Schwartz, and Flora M. Vaccarino. 2018. “Loss of TrkB Signaling in Parvalbumin-Expressing Basket Cells Results in Network Activity Disruption and Abnormal Behavior.” Cerebral Cortex 28 (10): 3399–3413. https://doi.org/10.1093/cercor/bhx173.

Yeung, Maggie S.Y., Sofia Zdunek, Olaf Bergmann, Samuel Bernard, Mehran Salehpour, Kanar Alkass, Shira Perl, et al. 2014. “Dynamics of Oligodendrocyte Generation and Myelination in the Human Brain.” Cell 159 (4): 766–74. https://doi.org/10.1016/j.cell.2014.10.011.

Yizhar, Ofer, Lief E. Fenno, Matthias Prigge, Franziska Schneider, Thomas J. Davidson, Daniel J. O’Shea, Vikaas S. Sohal, et al. 2011. “Neocortical Excitation/Inhibition Balance in Information Processing and Social Dysfunction.” Nature 477 (7363): 171–78. https://doi.org/10.1038/nature10360.

Zheng, Kang, Juan Ji An, Feng Yang, W. Xu, Z.-Q. D. Xu, Jianyoung Wu, T. G. M. Hokfelt, André Fisahn, Baoji Xu, and Bai Lu. 2011. “TrkB Signaling in Parvalbumin-Positive Interneurons Is Critical for Gamma-Band Network Synchronization in Hippocampus.” Proceedings of the National Academy of Sciences 108 (41): 17201–6. https://doi.org/10.1073/pnas.1114241108.

